# Combining MicroED and native mass spectrometry for structural discovery of enzyme-biosynthetic inhibitor complexes

**DOI:** 10.1101/2025.02.20.638743

**Authors:** Niko W. Vlahakis, Cameron W. Flowers, Mengting Liu, Matthew Agdanowski, Samuel Johnson, Jacob A. Summers, Catherine Keyser, Phoebe Russell, Samuel Rose, Julien Orlans, Nima Adhami, Yu Chen, Michael R. Sawaya, Shibom Basu, Daniele de Sanctis, Soichi Wakatsuki, Hosea M. Nelson, Joseph A. Loo, Yi Tang, Jose A. Rodriguez

## Abstract

With the goal of accelerating the discovery of small molecule-protein complexes, we leverage fast, low-dose, event based electron counting microcrystal electron diffraction (MicroED) data collection and native mass spectrometry. This approach resolves structures of the epoxide-based cysteine protease inhibitor, and natural product, E-64, and its biosynthetic analogs bound to the model cysteine protease, papain. The combined structural power of MicroED and the analytical capabilities of native mass spectrometry (ED-MS) allows assignment of papain structures bound to E-64-like ligands with data obtained from crystal slurries soaked with mixtures of known inhibitors, and crude biosynthetic reactions. ED-MS further discriminates the highest-affinity ligand soaked into microcrystals from a broad inhibitor cocktail, and identifies multiple similarly high-affinity ligands soaked into microcrystals simultaneously. This extends to libraries of printed ligands dispensed directly onto TEM grids and later soaked with papain microcrystal slurries. ED-MS identifies papain binding to its preferred natural products, by showing that two analogues of E-64 outcompete others in binding to papain crystals, and by detecting papain bound to E-64 and an analogue from crude biosynthetic reactions, without purification. This illustrates the utility of ED-MS for natural product ligand discovery and for structure-based screening of small molecule binders to macromolecular targets.

## Introduction

The development of new methods to interrogate miniscule protein crystals has enabled new frontiers for characterizing protein-ligand complexes. Laser and synchrotron-based X-ray sources continue to develop, delivering high resolution structures from assemblies of shrinkingly fewer repeat units (Holton & Frankel, 2010). This is facilitated in part by advances in serial crystallography at both synchrotrons and X-ray free electron lasers, which offer atomic resolution structures from slurries of nanocrystals (Colletier *et al*., 2016; Brewster *et al*., 2015; Roedig *et al*., 2017; Stellato *et al*., 2014). Complementing these efforts, microcrystal electron diffraction (MicroED), or 3D electron diffraction (3DED), yields high resolution crystal structures from necessarily thin (<500 nm) crystals, often capitalizing on a target’s proclivity to form small crystals (Shi *et al*., 2013; Nederlof *et al*., 2013; Yonekura *et al*., 2015), and enabling more rapid ligand diffusion and binding than is possible with large, X-ray diffraction (XRD)-suitable crystals (Clabbers *et al*., 2020; Martynowycz & Gonen, 2021).

The advantage of efficient ligand soaking in MicroED raises the possibility of rapid structural evaluation of ligand binding to proteins in high-throughput manner, especially where a binding site on the target protein is known. As such, a method that enables screening multiple candidate ligands against crystals of the protein of interest simultaneously, which can efficiently inform on if one or more has bound, would be an advantageous addition to the structural biology and drug discovery toolkit. This is illustrated by fragment-binding screens, where small molecule fragments representing a range of chemical space are simultaneously soaked into crystals of interest, typically prior to XRD (Chilingaryan *et al*., 2012; Erlanson *et al*., 2016; De Souza Neto *et al*., 2024). However, that approach is challenged when the bound ligand is not fully resolved or where resolution is insufficient for unambiguous atomic assignment (Verlinde *et al*., 2009; Patel *et al*., 2014). If the identity of a potential ligand is not known beforehand, a secondary validation method, such as mass spectrometry, can be useful (Cohen & Chait, 2001).

Native mass spectrometry (nMS) is ideal for this role since it allows for the observations of protein complexes and protein-ligand interactions in their near-native states, preserving non-covalent interactions (Tamara *et al*., 2022). Nano electrospray ionization (ESI) facilitates the de-solvation and ionization of proteins in a volatile buffer, and its coupling with mass spectrometry is capable of identifying species in complex mixtures with low sample volume (Fenn *et al*., 1989). In fact, nMS has played a crucial role in evaluating small molecule libraries for downstream applications in XRD soaking experiments (Gavriilidou *et al*., 2022), highlighting its potential for more high-throughput analysis. The application of nMS directly to samples used for MicroED (ED-MS), therefore offers the potential for accurate structures of protein-ligand crystals obtained from soaks with compound mixtures.

Here, we demonstrate the utility of the combined ED-MS approach for resolving protein-ligand complexes that are generated by soaking of protein microcrystal slurries with potential ligands. We use crystals of the cysteine protease papain as a scaffold on which to bind a panel of both known and novel potential inhibitors, including the epoxide-based covalent cysteine protease inhibitor E-64 (Varughese *et al*., 1989), its commercially available chemical analogues (Kim *et al*., 1992), and novel analogues generated by biosynthetic reaction (Liu *et al*., 2024). We determine structures of papain soaked with mixtures of these potential inhibitors, and perform nMS directly on TEM grid-adsorbed crystals following diffraction data collection to identify the exact masses of bound ligands present. Our results demonstrate that ED-MS can deliver high-quality structures of protein-ligand complexes directly from microcrystal slurries using tools readily accessible to a standard electron microscopy facility. The approach can further be leveraged to structurally illuminate the binding of proteins to novel natural products, and active components of inhibitor cocktails and biosynthetic reactions.

## Results

### 1. MicroED structures of papain bound to the cysteine protease inhibitor and natural product, E-64

Crystals of the cysteine protease papain (Fig. 1C), isolated from papaya latex, were grown and co-crystallized or soaked with target ligands for MicroED analysis. Papain crystals that diffracted to high resolution by X-ray diffraction (Fig. S1D,E), were readily crushed by repeated pipetting in their crystallization solution and frozen on cryoEM grids to yield a population of well-diffracting nanocrystalline fragments (Fig. 1A,B). Using samples prepared by this strategy, we acquired continuous rotation MicroED tilt series at 200 kV using a Direct Electron Apollo detector and determined a 2.5 Å resolution structure of papain phased by molecular replacement (Kamphuis *et al*., 1984) (Fig. 1E, Table S1).

**Figure 1.**
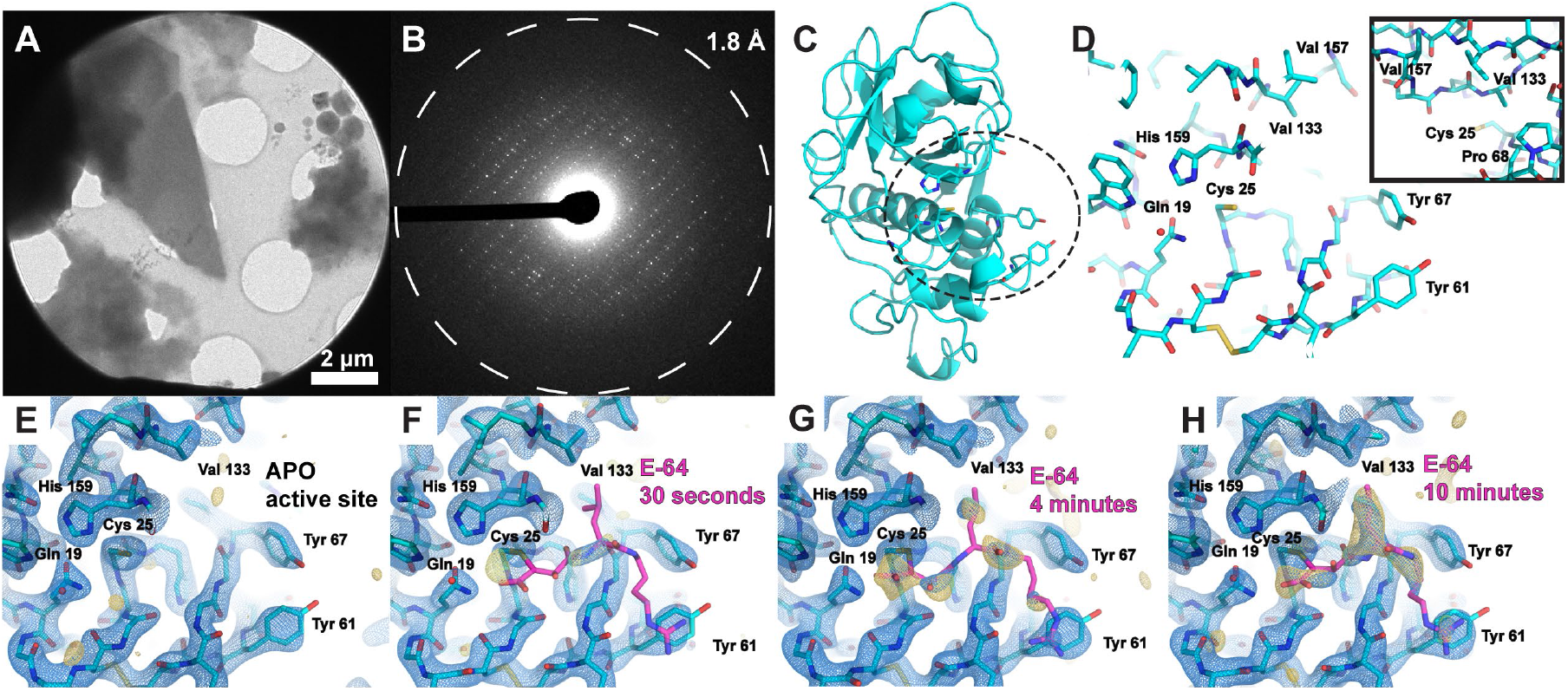
MicroED structures of papain and E-64-papain complexes. TEM image of a thin frozen-hydrated papain microcrystalline fragment (A), which yields high resolution electron diffraction when illuminated with a low-flux parallel beam (B). Atomic model of the cysteine protease papain, determined by MicroED to 2.5 Å resolution with secondary structure and key active-site residue side chains rendered, with dashed circle highlighting the active site (C). View of papain’s active site from the same model, with key residues labeled. Covalent inhibitors bind in this pocket to cysteine 25, and a hydrophobic pocket (view in inset) constituting valine 133, valine 157, and proline 68 accommodates aliphatic moieties of potential binders (D). Structure of the apo-form of papain at 2.5 Å resolution determined by MicroED from crystals such as the one shown in panel A, where the active site is viewed (E). MicroED structures of the papain active site at 2.5 Å resolution from microcrystals soaked with a concentrated solution of E-64 for 30 seconds (F), 4 minutes (G), and 10 minutes (H). Blue mesh indicates the 2F_o_-F_c_ map at 1.5σ level following modeling of the ligand (magenta coordinates), and gold mesh indicating the F_o_-F_c_ map at 3σ level that was present prior to modeling a ligand. 10 minutes of soaking is sufficient for high (>80%) occupancy to be achieved.

We next validated that the known epoxide-based cysteine protease inhibitor E-64 covalently bound papain (Fig. S2A,B) and could be detected by MicroED when soaked into papain microcrystal slurries. Given the potential speed advantage in ligand soaking with the thin crystals used for MicroED, we characterized how quickly the measured occupancy of E-64 increased in the active sites of papain microcrystals as a function of soaking time. Papain microcrystals frozen on cryoEM grids following either 30 seconds, 4 minutes, or 10 minutes of soaking in 2.5 mM E-64 were interrogated by MicroED, each yielding a structure with a clear view of the binding pocket surrounding critical residue cysteine 25 at 2.5 Å resolution (Fig. 1F,G,H, Table S2). Only a weak trace of density indicating E-64 was observed following 30 seconds of soaking (Fig. 1F), but for both the 4 and 10 minute structures sufficient density was present to model E-64 and carry out refinement (Fig. 1G,H). The soaking efficiency noted in microcrystals exceeded that of equivalently soaked macroscopic crystals needed for single-crystal XRD (Fig. S6, Table S7).

Although achieving high completeness MicroED data can require the merging of partial datasets from multiple crystals, we noted that the clearest density in papain’s active site in F_o_-F_c_ ligand omit maps was often achieved from unmerged or minimally merged datasets, provided completeness exceeded approximately 85%. This is likely due to variable ligand occupancy between different microcrystals on the same TEM grid (Fig. S3). As such, the ideal data collection modality required wedges of data that are as large as possible, to be quickly acquired on individual papain crystals prior to much accumulation of radiation damage. This was uniquely facilitated by the use of a fast direct electron detector (DE Apollo) performing event based electron counting (Hattne *et al*., 2023; Vlahakis *et al*., 2024). Leveraging the DE Apollo camera for fast, low-flux MicroED data collection (Peng *et al*., 2023), we were able to routinely acquire structure-worthy tilt series delivering no more than 6 e^-^/Å^2^ total fluence per crystal.

### 2. Unambiguous determination of papain-E-64-like natural product complexes enabled by ED-MS

We anticipated MicroED would be a useful tool for ligand discovery from novel natural product libraries. We therefore proceeded to determine the structure of papain microcrystals co-crystallized in the presence of an E-64 analog recently discovered by novel combinatorial biosynthesis platform (Liu *et al*., 2024), which we refer to as E-64-A65 (Fig. S2C,D). A 2.5 Å resolution structure of the papain-E-64-A65 cocrystal complex showed prominent density in the active site, confirming the bound biosynthetic compound (Fig. 2F, Table S1). However in resolving this structure a potential limitation of the approach was noted: although MicroED was able to rapidly deliver a structure of the bound ligand complex, the ligand density was not always sufficient to disambiguate between structurally similar binders such as E-64-A65 and its parent, E-64. This is evident by comparing the E-64-A65-bound structure to a high-occupancy structure of papain co-crystallized with E-64 and determined to 2.3 Å resolution (Fig. 2D, Table S1). This scenario would be common during screening of crystal binding to highly similar ligands, or to mixtures or cocktails of similar ligands, as might be performed in high-throughput drug discovery efforts. As such, we turned to native mass spectrometry (nMS), performed on crystals dissolved from the same cryoEM grid, following MicroED data collection (Fig. 2A, Fig. S4).

**Figure 2.**
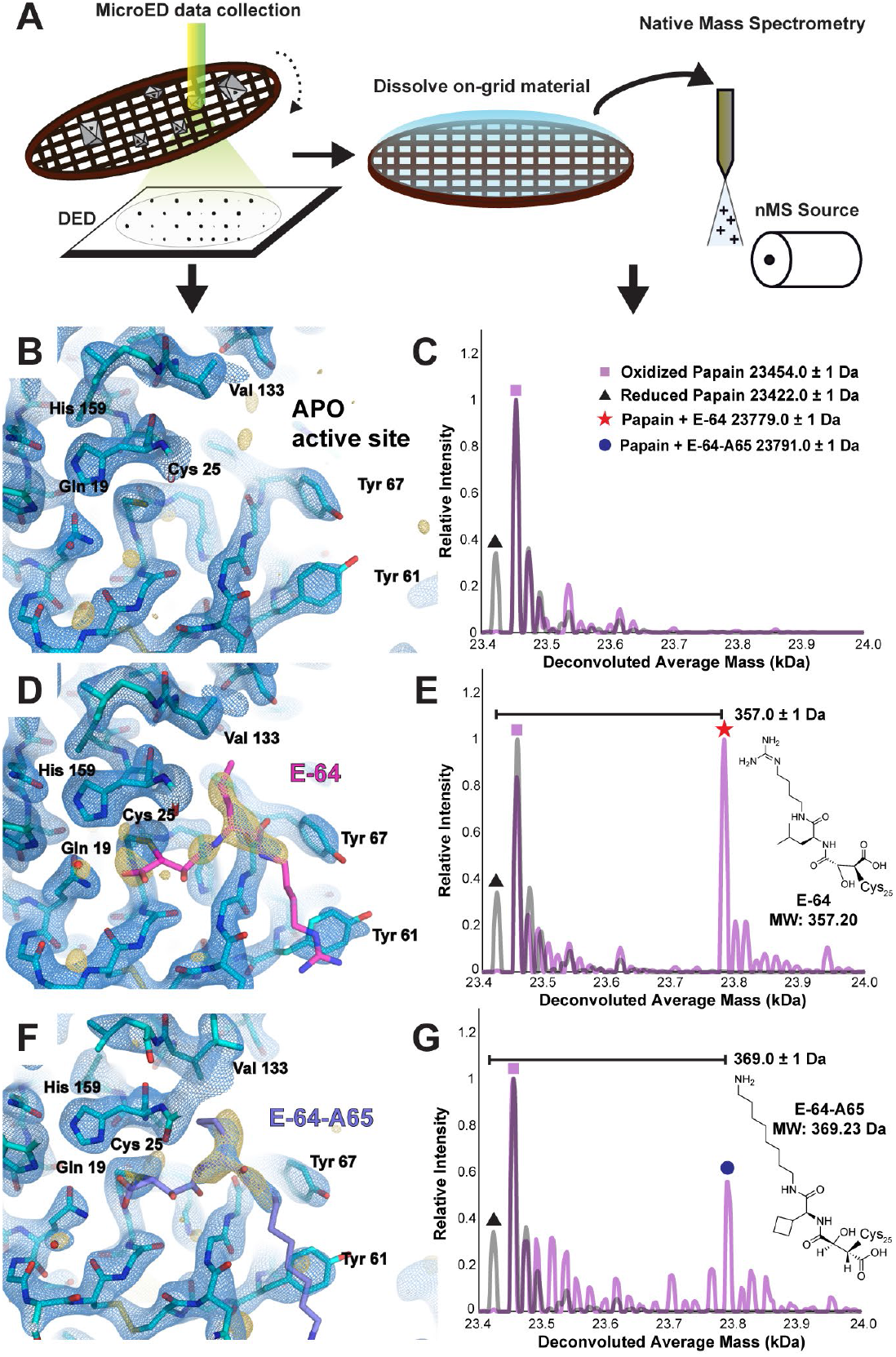
Native mass spectrometry of MicroED samples confirms ligand identity. Cartoon schematic detailing process of collecting MicroED data from ligand-bound crystals on a direct electron detector (DED), then recovering the sample and dissolving the on-grid material for nMS analysis (A). Structure of the apo-form of papain at 2.5 Å resolution determined by MicroED from crystals such as the one shown in panel C, where the active site is viewed (B), and nMS spectrum of the same material harvested and dissolved from the cryoEM grid (C). Structure of the papain-E-64 co-crystal complex at 2.3 Å resolution determined by MicroED (D), and corresponding nMS spectrum confirming presence of the papain-E-64 complex on the sample (E). Structure of the papain-E-64-A65 co-crystal complex at 2.5 Å resolution determined by MicroED (F), and corresponding nMS spectrum confirming presence of the papain-E-64-A65 complex on the sample (G). Blue mesh indicates the 2F_o_-F_c_ map at 1.5σ level following modeling of the ligand, and gold mesh indicates the F_o_-F_c_ map at 3σ level that was present prior to modeling a ligand.

Mass spectra acquired from apo papain crystals harvested from a TEM grid showed masses corresponding to the protein with both a doubly oxidized, sulfinic acid form (+32 Da) of its catalytic residue Cys25, and a reduced form of the protein (Fig. 2C). The assignment of this oxidation state is substantiated by its resistance to reduction with DTT or TCEP, unlike other +32 Da modifications such as persulfides, which would be reduced under these conditions (Fig. S5). By comparison, mass spectra acquired from on-grid material of both papain-E-64 and papain-E-64-A65 cocrystals offered unambiguous confirmation of the identity of each bound species. A 357 Da and 369 Da mass shifted species relative to the reduced form of papain, in the mass spectrum acquired from papain-E-64 and papain-E-64-A65 samples respectively, dominated in each, matching the expected molecular weights of the ligands: E-64 (357.20 Da) and E-64-A65 (369.23 Da) (Fig. 2E,G, Table S13). These results therefore validated ED-MS as an effective approach for validating the mass of bound compounds within microcrystals used for structure determination, informing on both the identity and bound structure of an unknown ligand.

#### 3. Structure of papain-E-64 captured by ED-MS from microcrystal slurry soaked with an inhibitor cocktail

Having validated that we could quickly soak ligands into papain microcrystals, visualize them by MicroED, and confirm their identities by nMS, we verified the utility of this workflow for higher throughput ligand discovery efforts. In particular, we aimed to test if the same information about ligand presence, conformation, and identity could be extracted by ED-MS for papain crystals soaked with a mixture of potential ligands. To evaluate this, we soaked papain microcrystals with a commercially available protease inhibitor cocktail containing E-64, AEBSF, leupeptin, aprotinin, bestatin, and EDTA at varying relative concentrations (Fig. 3A, Fig. S9). Of these components, both E-64 and leupeptin are known ligands of papain. Cocktail-soaked papain microcrystals were then deposited and plunge-frozen on TEM grids and interrogated by ED-MS (Fig. 3B). A 2.5 Å resolution MicroED structure revealed unsatisfied active-site density in the absence of a modeled ligand (Fig. 3C). nMS analysis confirmed that the papain-E-64 complex was the only bound species dominantly present in the TEM grid-adsorbed crystals (Fig. 3D). Therefore, E-64 was subsequently modeled into the unassigned density in the final steps of structure refinement (Fig. 3E, Table S3). Even with the improved resolution afforded by single crystal XRD, distinguishing E-64 from leupeptin based solely on the appearance of active site density in a 1.5 Å structure from macroscopic crystals soaked with the same cocktail (Table S8) was non-trivial without the additional information afforded by nMS (Fig. S8).

**Figure 3.**
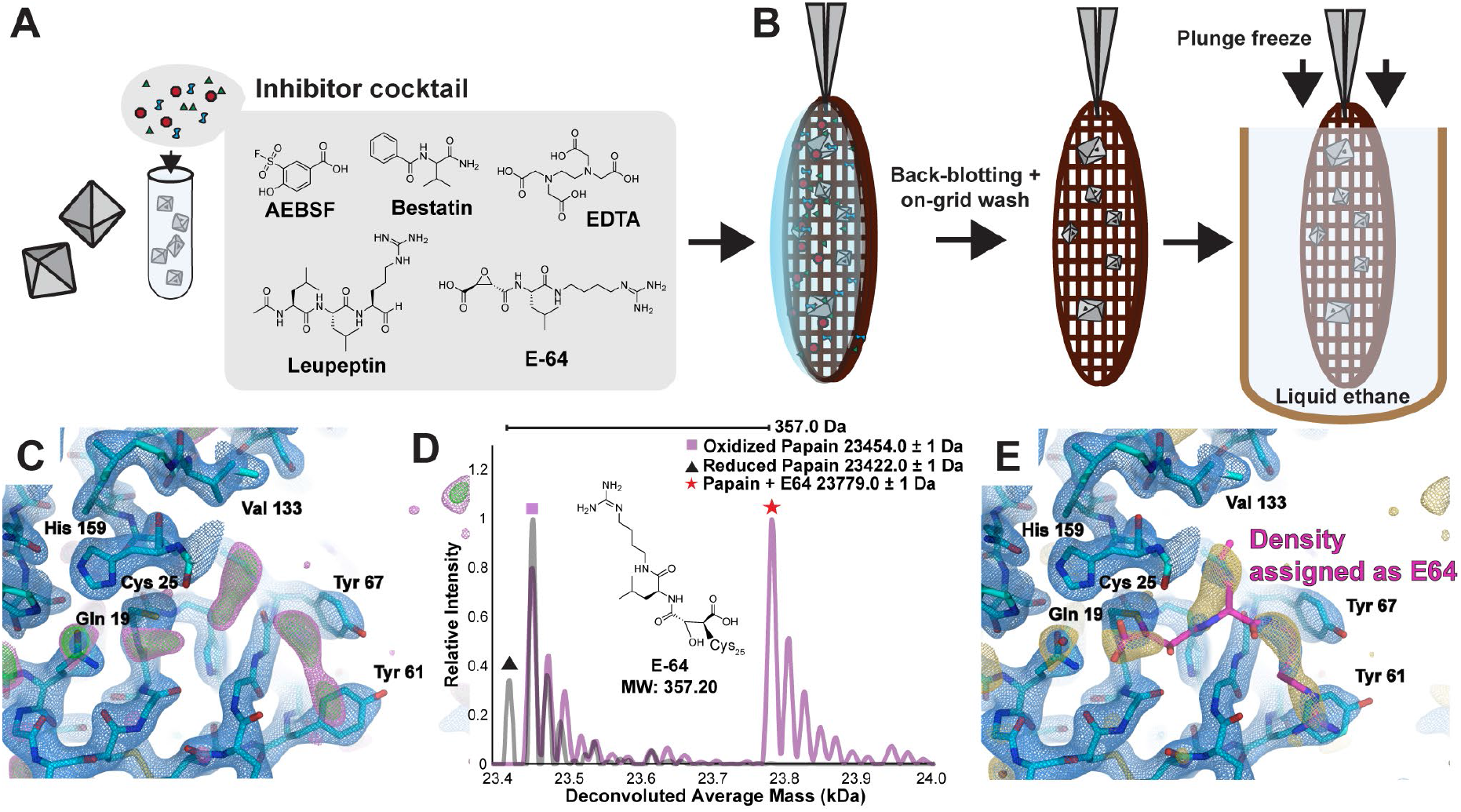
ED-MS reveals the structure of the papain-E-64 complex from microcrystals soaked with protease inhibitor cocktail. Papain crystals were crushed in suspension and soaked with a commercial inhibitor cocktail solution containing AEBSF, bestatin, EDTA, leupeptin, aprotinin, and E-64 (A). Microcrystalline suspension was loaded on a holey carbon cryoEM grid, back-blotted, washed with compatible solvent, then back-blotted away prior to plunge freezing (B). MicroED data from this sample yielded a 2.4 Å resolution structure with clear difference map density occupying the papain active site (F_o_-F_c_ map displayed at 2.5σ level in magenta and 3σ level in green) (C). nMS spectrum of the on-grid material reveals presence of the papain-E-64 complex and no other bound species (D). The revised model of papain’s active site with the bound inhibitor E-64 modeled in is shown in panel E, with the 2.5σ F_o_-F_c_ ligand-omit map overlayed in gold.

Notably, mixing of solubilized papain with leupeptin alone and measuring nMS of the solution revealed that leupeptin does indeed bind papain (Fig S10B). Likewise, when crystallized in the presence of a molar excess of leupeptin, single-crystal X-ray diffraction shows unambiguous density for bound leupeptin in the papain active site (Fig. S10A, Table S9). Therefore, the results of the inhibitor cocktail soaking experiment indicate that, if multiple binders are present in the soaking mixture, the higher affinity binder may outcompete the others and be unambiguously identified by ED-MS at the short time and length scales evaluated.

### 4. ED-MS applied to papain bound to mixtures of its high affinity ligands, E-64, E-64C and E-64D

We next aimed to test the limits of structural information achievable by ED-MS for crystals soaked with mixtures of multiple high affinity binders. In addition to E-64, papain is known to bind its commercially available analogs E-64C and E-64D (Kim *et al*., 1992), and their complexes have known structures (Liu et al. in press, PDB IDs: 9CLH, 9CKT, 9CKW, 9EG7). Given the results of ED-MS analysis on inhibitor cocktail-soaked crystals, we expected a similar experiment might reveal whether one of these known high-affinity ligands could outcompete the others when mixed in equimolar quantity and soaked into papain microcrystals. A 2.5 Å resolution MicroED structure was determined from mixture-soaked papain crystals, where unsatisfied density in the active site was detected that could not be confidently assigned to any one of the three structurally similar candidate compounds (Fig. 4A). nMS of the same crystals revealed the mass shifts corresponding to the E-64 and E-64C bound states of papain were both present in the mixture (Fig. 4B). The active site of the MicroED structure was therefore modeled as an ensemble of the papain-E-64 and papain-E-64C bound states (Fig. 4C, Table S3). Even with a higher resolution structure of papain microcrystals soaked with the same mixture determined by serial synchrotron X-ray diffraction, the residual density in the active site only clearly showed the structural fragment common to all three ligands (Fig. S16, Table S10).

**Figure 4.**
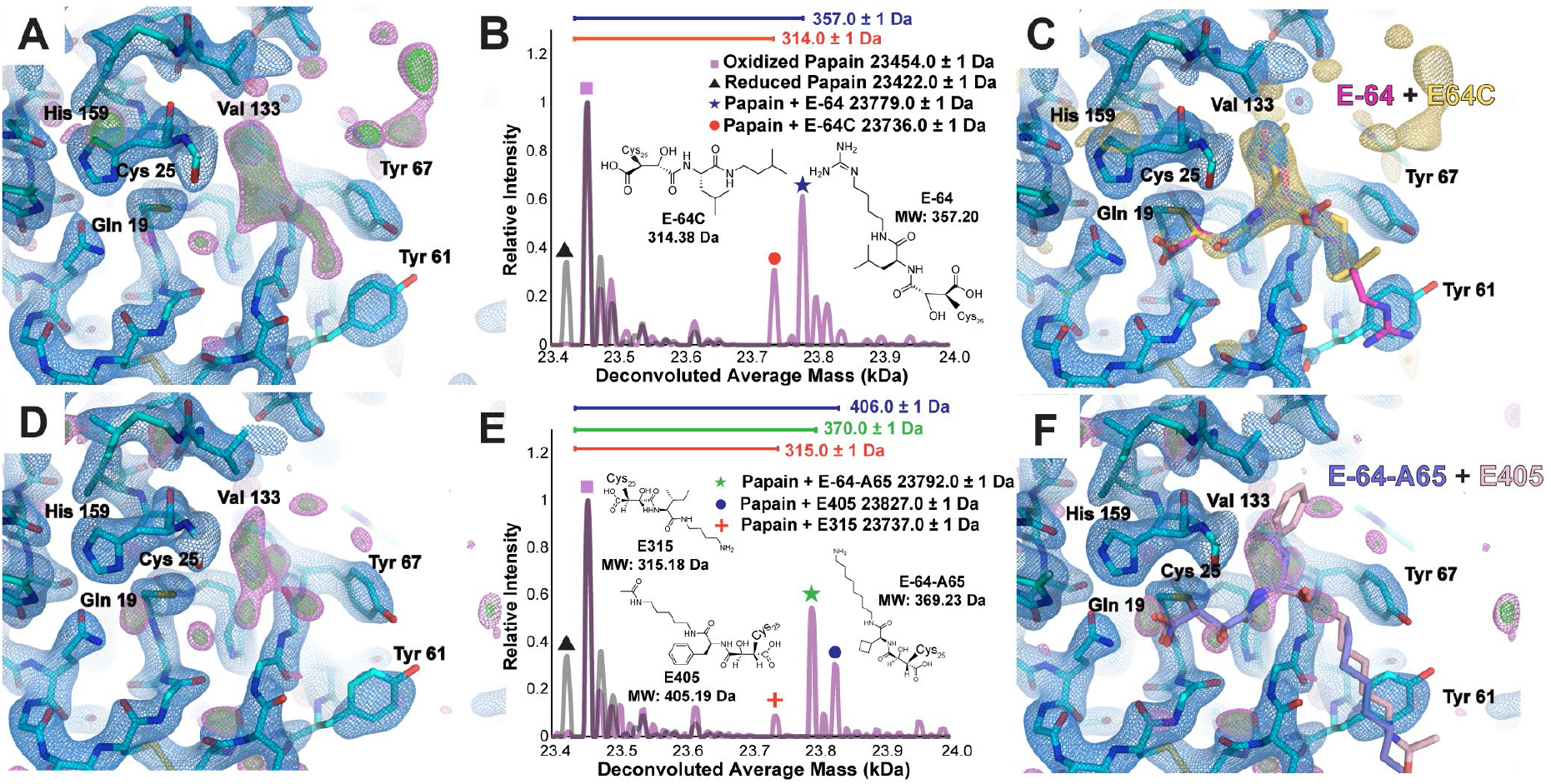
Complex structures from papain microcrystals soaked with mixtures of known and unknown commercial and novel biosynthetic binders. Active-site view of the MicroED structure of papain microcrystals soaked with an equimolar mixture of the high-affinity inhibitors E-64, E-64C, and E-64D at 2.5 Å resolution, with unassigned density in the difference Fourier map (F_o_-F_c_ map displayed at 2.5σ level in magenta and 3σ level in green, panel A). nMS spectrum of the on-grid material from this sample showing presence of the papain-E-64 and papain-E-64C complexes (B), and fully refined active site visualization with E-64 and E-64C modeled as alternate conformations in the MicroED structure with the 2.5σ F_o_-F_c_ ligand-omit map overlayed in gold (C). Active-site view of the MicroED structure of papain microcrystals soaked with an equimolar mixture of novel biosynthetic E-64 analogues E-64-A65, E315, E371, and E405 at 2.4 Å resolution with unassigned difference map density (D). The nMS spectrum from the grid that yielded this structure shows presence of primarily the papain-E-64-A65 complex and papain-E405 complex, and trace amounts of the papain-E316 complex simultaneously (E). While the ligand density was not sufficiently clear to enable accurate modeling of these ligands in the binding pocket, the structures of E-64-A65 and E405 in their conformations determined by X-ray diffraction are shown overlayed onto the unsatisfied density (F).

Curiously, when interrogating papain-E-64C and papain-E-64D co-crystal complexes separately by ED-MS (Fig. S11, Table S5), nMS revealed in each case that only the bound species corresponding to E-64C, indicated by a 314 Da mass shift from the reduced apo-form papain species, was present (Fig. S11C,F). To confirm the veracity of the stock E-64D, we determined its structure directly from the microcrystalline powder used for the ligand soaking experiment, and noted the refined 0.8 Å resolution structure (Fig. S12G, Table S12) matched the expected structure of E-64D, with a molecular weight of 342 Da. Likewise, LCMS of the E-64D dissolved in 66% methanol solution revealed the 342 Da mass (Fig. S12C,D). The only structural difference between E-64C and E-64D is the presence of an ester, rather than a terminal carboxylic acid, bound to the epoxy warhead. We therefore concluded that that the ester of E-64D is likely hydrolyzed specifically in the course of its covalent attachment to the papain active site cysteine, effectively converting into E-64-C, as observed in the crystal structure. This hypothesis is consistent with previous work studying the increased likelihood of ester hydrolysis in the presence of neighboring sulfides (Rydholm *et al*., 2007).

### 5. ED-MS of papain bound to on-grid arrayed ligand sets

Given the ability of ED-MS to identify papain complexed to multiple known, high affinity binders when mixed and used as a cocktail to soak protein microcrystals, we next applied the method to interrogate papain crystals applied to grids on which one or more target ligands had been pre-arrayed. We reasoned that the printing of potential inhibitors on grids using microarrayer devices would streamline high throughput soaking and ED-MS efforts. The arrayED technique (Delgadillo *et al*., 2024) was used to deposit 144 250-350 picoliter droplets of E-64 and leupeptin onto TEM grids, which were then dried and stored until ready for soaking. A slurry of papain microcrystals was then applied to the arrayED grids and incubated for 5 minutes before freezing and ED-MS analysis. Data collected from these grids yielded a 2.8 Å structure of papain complexed with E-64 (Fig. S7E, Table S6), and nMS on the same grid revealed distinct presence of the papain-E-64 complex on the sample (Fig. S7F).

### 6. ED-MS detects high affinity papain-inhibitor complexes from a cocktail of biosynthetic E-64 derivatives

To test the utility of ED-MS for drug discovery efforts from mixtures of natural product libraries, we soaked papain microcrystals with an equimolar mixture of multiple novel biosynthetic E-64 analogs. This collection included E-64-A65, as well as a series of similar compounds generated by substituting different amino acids and amine groups as precursors for the biosynthetic reaction catalyzed by Cp1B and Cp1D. These compounds, with molecular weights of 315, 371, and 405 Da, are referred to as E315, E371, and E405 (Fig. S14). While nMS of on-grid crystals soaked in a mix of these compounds found clear signal for both papain-E-64-A65 and papain-E405 complexes, and trace signals for the papain-E315 complex, only weak density could be observed in the active site of a 2.4 Å MicroED structure of the mixture (Fig. 4D,E, Table S3). Nonetheless, omit difference maps showed residual active site density that partially matched the more ordered fragments of E-64-A65 and E405 (Fig. 4F). Given the challenge in refinement of the mixed structure model, excluding the strongest binders from otherwise equivalent mixtures could help clarify the ambiguous density and discover lower affinity ligands. Repeating the same experiment, but excluding E-64-A65 from the cocktail, we measured masses for papain complexes with each of the three other natural products, though active site density from the MicroED structure of these crystals was even weaker in the absence of the highest-affinity ligand (Fig. S13, Table S3). Serial synchrotron X-ray crystallographic structures of papain microcrystals soaked with each of the individual components of the mixture, E-64-A65, E315, E371 and E405, support these ED-MS results, where active site density in these structures confirmed E405 as a clear binder to papain (Fig. S17D, Table S11).

### 7. Determination of papain binders directly from crude biosynthetic reaction mixtures by ED-MS

Isolation of novel natural products is often non-trivial, and achieving comparable yields to commercially available compounds can be limiting. In such cases, we anticipate broad utility of ED-MS for discovery of binders from heterogeneous research samples, natural product extracts, and reactions performed as part of drug discovery efforts, despite the potentially low concentration of binders in those samples. To test this use-case, we soaked papain microcrystals directly with biosynthetic reaction mixtures, where E-64 and E-64-A65 were each the intended major products (Fig. 5A,B,C). Still containing reaction precursors, side products, and enzyme catalysts, the crude solutions contained at most 1 mM of the intended E-64 product, and 2 mM of the E-64-A65 product (Fig. S15). To compensate for this limited concentration, microcrystals were allowed to soak for a longer period of 4 hours prior to freezing on cryoEM grids. MicroED data collection on these microcrystals yielded a 2.3 Å resolution structure of the crude E-64-soaked crystals (Fig. 5D, Table S4), and a 2.5 Å resolution structure of the crude E-64-A65-soaked crystals (Fig. 5F, Table S4). In each structure, prominent density in the active site indicated the presence of a bound inhibitor. nMS of the on-grid material once again revealed that the mass of the ligand-bound complex matched the papain-E-64 complex, for the crude E-64 soaking trial (Fig. 5E), and the papain-E-64-A65 complex, for the crude E-64-A65 trial (Fig. 5G). In neither experiment was a species corresponding to papain bound with a precursor or side product of the biosynthetic reaction detected. This indicates the promise of ED-MS for natural product-based drug discovery efforts, where discovered compounds are available only in limited concentration or purity. It further suggests a potential application to high-throughput ligand screening by directly soaking crude reaction mixtures into microcrystals.

**Figure 5.**
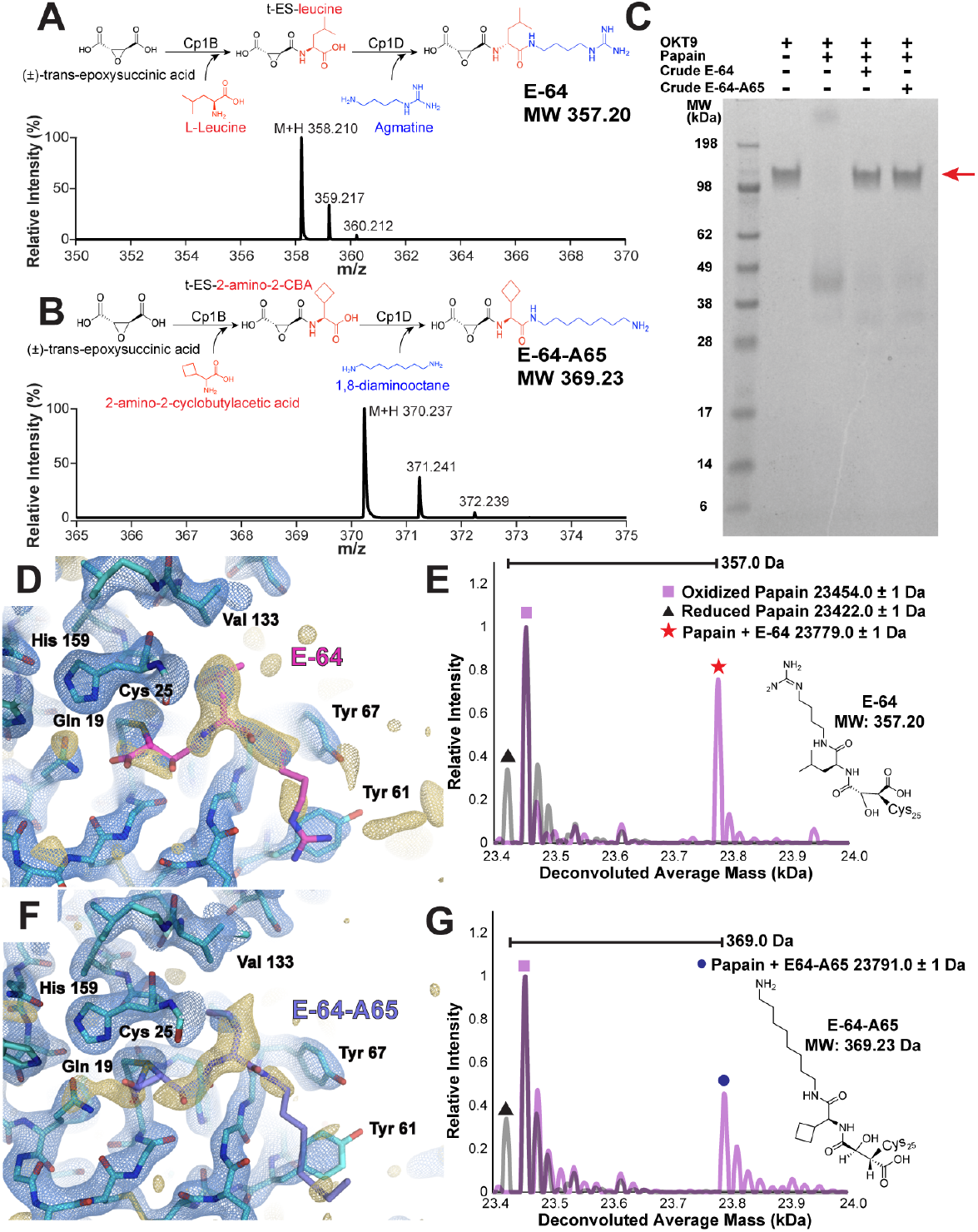
Strategy and demonstration of structural characterization of protein microcrystals soaked with crude biosynthetic products by ED-MS. Schematic for biosynthetic production of E-64 (A) and E-64-A65 inhibitors (B) with corresponding LC-MS traces. SDS-PAGE demonstrating activity of papain in cleaving the 150 kDa antibody OKT9, where when only papain and OKT9 are present the 150 kDa band disappears indicating successful cleavage. When papain is pre-treated with either crude preparation of biosynthetic products E-64 or E-64-A65, the 150 kDa band for OKT9 is retained indicating successful inhibition of the protease (C). MicroED structure of papain microcrystals soaked with this preparation of E-64 without additional purification, showing the active site at 2.3 Å resolution with density indicating the presence of the ligand (D), and nMS spectrum from this sample indicating the presence of the papain-E-64 complex (E). MicroED structure of papain microcrystals soaked with a similar crude E-64-A65 preparation without additional purification, showing the active site at 2.5 Å resolution (F), and nMS spectrum from this sample indicating the presence of the papain-E-64 complex (G). Blue mesh indicates the 2F_o_-F_c_ map at 1.5σ level following modeling of the ligand, and gold mesh indicates the F_o_-F_c_ map at 2,5σ level that was present prior to modeling a ligand.

## Discussion

Drug discovery efforts rely on X-ray crystallography as a gold standard method for structural analysis of protein target-ligand complexes (Chilingaryan *et al*., 2012). These techniques excel when large crystals of high quality can be grown, or when large volumes of concentrated crystal slurries are available. However, for microcrystalline samples, or where crystal quantity or yield may be limited, electron diffraction promises to be a valuable addition to this toolkit. MicroED has proven robust at delivering molecular-resolution structures of protein micro- and nanocrystals, and has been anecdotally applied to the determination of protein-small molecule complexes. However, the application of MicroED to more systematic screening of bound drug-like ligands has yet to be thoroughly evaluated. Also understudied are the limitations of MicroED in this context, and the strategies for overcoming them to deliver accurate, unambiguous structural information about protein-ligand complexes.

We find that MicroED can be readily leveraged to determine accurate, molecular resolution structures of well-ordered protease microcrystals co-crystallized and soaked with a panel of potential ligands. Structures of papain-ligand complexes that afforded the most distinct density in the electrostatic potential map for bound ligands were determined from tilt series acquired on single microcrystals. Each crystal exposed to less than 4.5-6.0 electrons per square Angstrom total fluence using 200 keV electrons, mitigating beam-induced damage (Vlahakis *et al*., 2024). Damage plays a role in data quality, since structures determined from data within this dose budget showed evidence of radiation damage, including broken disulfide bonds. In this context, accurate detection of weak diffraction signal is impactful. To limit dose, but record high completeness data, we used a fast, sensitive direct electron counting detector (Vlahakis *et al*., 2024). Under these conditions, merging of partial datasets benefits completeness, but in turn reduces the clarity of ligand signal in the active site due to crystal non-isomorphism and variable occupancy between crystals. Although rare, the optimal condition is achieved when sufficiently complete data are collected from a single crystal, or when uniform occupancy is achieved across crystals. Alternative strategies for limiting the radiation damage delivered to microcrystals during diffraction acquisition have enabled clear resolution of bound ligands. Notably, leveraging the TEM’s nanoprobe mode to illuminate multiple distinct μm areas of single crystals during tilt series of narrow angular size yielded high data completeness from only two crystals while mitigating the impact of radiation damage on the data, and ultimately revealing distinct density for a bound peptidic ligand (Shaikhqasem *et al*., 2025).

While MicroED delivered ready confirmation of a ligand’s presence in a target binding site, the observed E-64 density is often partial, lacking clarity at its more flexible, solvent exposed terminus. For its analogues and derivatives, this challenged unambiguous *de novo* identification. This limitation is not an issue when protein crystals are soaked with a solution of only one compound of interest, but becomes critical when they are soaked with mixtures; the latter are important for high-throughput screens, or when investigating less pure research samples. We find that complementing MicroED with nMS of grid-adsorbed crystals, following MicroED data collection, provides an efficient solution to this problem and enables unambiguous verification of which bound species are present in a sample of mixture-soaked microcrystals. In addition to reliably validating the identity of ligands present in structures from controlled soaking and co-crystallization experiments on papain and E-64 analogs, ED-MS was able to inform on papain complexed with mixtures of potential binders that MicroED alone may not have resolved. Application of creative data collection strategies for limiting the impact of radiation damage on MicroED data might improve the likelihood of *de novo* ligand identification (Shaikhqasem *et al*., 2025) in cases where nMS is not available as a complementary method.

While ED-MS can be scaled as a high-throughput solution for fragment-based drug discovery, some challenges remain. Automated routines for cryogenic MicroED data collection are still limiting, when compared to those routinely applied to CryoEM single particle and tomography acquisition (De La Cruz, 2021; Bücker *et al*., 2020). As these methods advance, we anticipate that the efficiency of data acquisition for ED-MS efforts will starkly improve and allow for more broad screening of potential ligands to a target than was achievable here. Additionally, depending on the identity of the target protein crystals, particular optimization of sample preparation conditions might often be necessary, as common crystallization reagents such as polyethylene glycol may confound mass spectrometry analyses of grid-adsorbed material (Zhao & O’Connor, 2007). Such optimization efforts might benefit from ligand printing on grids at scale, facilitated by arrayED technologies (Delgadillo *et al*., 2024), and from advances to sample preparation approaches that facilitate nMS data collection.

By allowing for the determination of ligand-bound protein structures directly from biosynthetic reaction mixtures, ED-MS shows promise for natural product-focused ligand discovery efforts, where compounds are often of limited concentration and purity. This workflow can also be readily applied to automated ligand fragment screening, mimicking experimental schemes increasingly used in synchrotron and XFEL experiments but foregoing the requirements of rare beamtime and sufficiently large crystals. In such a case, MicroED would offer a first indication of binders present in a given mixture, and would provide an essential picture of any binding sites detected on the target protein, while nMS verifies precisely which fragments or components of the mixture successfully bound. Finally, the application of nMS to TEM grid-adsorbed samples promises to be readily translatable to other cryoEM modalities, indicating potential for similarly correlative measurements to deliver unique insights in single-particle imaging and tomography experiments.

## Materials and Methods

### Crystallization and MicroED sample preparation

Twice-crystallized papain extract from papaya latex in buffered aqueous suspension at approximately 25 mg/mL protein concentration was purchased from Sigma and used for crystallization experiments without further purification. Crystals formed by vapor diffusion in sitting drop trays, where the commercial papain suspension was mixed with methanol at a 1:2 volume ratio of papain to methanol. A volume of 15 μL was typically targeted in each well, incubated with a reservoir solution containing 59% methanol and 889 mM NaCl. Large single crystals suitable for single-crystal XRD would typically grow within 48 hours under these conditions. Such crystals were resuspended in 5-10 μL of reservoir solution, crushed by repeated pipetting, and used to seed fresh crystals by transferring small amounts of crystal fragments with horsehair and streaking across a freshly prepared sitting drop prior to incubation. For co-crystallization experiments, each potential ligand was dissolved in 66% methanol and included in the crystallization well, such that it was present at 1.6 times molar excess of papain in the well.

Crystals were resuspended in their sitting drops in a solution of 59% (v/v) methanol and 296 mM NaCl, and crushed by repeated pipetting to produce a 5 μL slurry of microcrystalline fragments. For structure determination of apo-form papain and ligand-soaked papain crystals, 0.5 μL of 500 mM DTT was added to the slurry. For each ligand soaking experiment, 4 μL of ligand solution (5.1 mM pure E-64, or between 1-2 mM crude E-64 or E65-A65) was added to the crystal slurry a set amount of time prior to plunge freezing. For pure E-64, samples were frozen 30 seconds, 4 minutes, and 10 minutes after the ligand was added. For microcrystals soaked with protease inhibitor cocktail, samples were frozen both 10 minutes and 4 hours after adding the cocktail. For microcrystals soaked with crude E-64 and crude E-64-A65, samples were frozen four hours after ligand was added to the slurry. For papain-ligand cocrystals, no additional treatment was delivered after crushing crystals into slurries prior to freezing.

All cryoEM samples were prepared on R2/1 Quantifoil holey carbon grids with a copper mesh. Grids were glow discharged using a PELCO easiGlow system on each side for 30 seconds each, with a plasma current of 15 mA, then mounted in an FEI Vitrobot with a humidity-saturated chamber held at 12 oC. 1 μL of 66% methanol was loaded on the back side of each grid, and 1.5 μL crystal slurry was loaded on the front. Excess solvent was blotted away from the grid by manually contacting only the back side with a piece of filter paper gripped in a long pair of tweezers for 10 seconds. An additional 1 μL of 66% (v/v) methanol was then quickly added to the same side of the grid crystals had been applied to wash away excess salt from the mother liquor and immediately blotted away, once again by blotting only the back side of the grid. The grid was then plunge frozen in liquid ethane and stored under liquid nitrogen until transferred to the TEM.

### MicroED data acquisition

MicroED data was collected at a ThermoFisher Scientific Talos F200C TEM operating at 200 keV, equipped with both a Direct Electron Apollo detector with a native frame rate of 60 Hz (Peng *et al*., 2023), and a Ceta-D scintillator-based camera. A side-entry cryotransfer holder was used to preserve each grid used for data collection at 100 K. For each sample, a grid atlas was first acquired at 155x magnification on the DE Apollo using SerialEM. Promising thin crystals with sharp edges were identified from these maps and navigated to. These crystals were aligned to eucentricity at higher magnification using imaging-mode settings at 4300x magnification, spot size 11, with the C2 lens condensed to ~44.8 %. This configuration gave an electron beam flux density of approximately 0.01 e^-^/Å^2^s on the sample, which was suitable for minimizing dose delivered to each crystal prior to diffraction data collection. Diffraction was measured by centering each crystal of interest within a 100 μm diameter selected area aperture projecting an area approximately 3 μm in diameter onto the specimen, and illuminating the crystal in diffraction mode with a parallel beam, configured at spot size 10 with a C2 lens current of ~45.5% and a virtual camera length of 960 mm. This configuration produced an electron beam flux density of approximately 0.03 e^-^/Å^2^s on the sample. A single 3 second diffraction exposure was acquired on each crystal using the Ceta-D detector to assess if it diffracted well, while subsequent acquisition was performed on the DE Apollo. Continuous-rotation tilt series in diffraction mode were acquired on each crystal targeting as wide an angular wedge as was achievable without the crystal of interest becoming obstructed, typically no more than 120o. The stage rotation speed was set to 0.6 o/s, paired with 90-frame integration on the DE Apollo, to produce a movie of images corresponding to 1.5 second effective exposures and 0.9 o/frame oscillations through reciprocal space. An alternative modality, where the stage rotation speed was set to 0.3 o/s paired with 180-frame integration on the DE Apollo, was occasionally used for more weakly diffracting crystals, where allowing greater electron flux per final frame of the movie appeared to positively influence the measurable diffraction signal. However, this slower configuration typically necessitated the merging of more partial datasets to achieve high data completeness, as crystals would suffer from radiation damage prior to completing a tilt series. In general, the faster acquisition strategy was more ideal for recording high-completeness datasets with more favorable data reduction statistics. All movies were collected using a binning factor of 2 during acquisition, such that frames were 4096×4096 pixels in size. Following MicroED data collection, grids were recovered into liquid nitrogen and stored until they could be analyzed by nMS.

### MicroED data processing and structure determination

All MicroED movies from the DE Apollo camera were saved in MRC file format and converted to SMV image stacks using a script developed in-house and run using MATLAB version 2023b. The only modification made to pixel values in these movies during conversion was a universal addition of 1 to all pixels, such that no counts of zero were present in the converted images. Indexing and data reduction was performed using *XDS* (Kabsch, 2010*b*), and scaling and merging were performed using *XSCALE* (Kabsch, 2010*a*). For each dataset, phase retrieval was achieved by molecular replacement with a known model of apo-form papain (PDB ID: 9PAP) (Kamphuis *et al*., 1984) using *PHASER* (McCoy *et al*., 2007). Structure refinement was performed in *PHENIX* (Adams *et al*., 2010) by the following strategy: reference model restraints from the original search model from the PDB were applied, while three cycles of refinement of XYZ coordinates in real and reciprocal space, group B-factors, and occupancies were performed. The model and map were then examined in *COOT*, and the model was edited to better agree with the measured data as indicated by maps in the F_o_-F_c_ map. Water molecules were modeled in where positive peaks in the F_o_-F_c_ map exceeding 3 sigma levels were found, if a hydrogen-bonding partner between 2.5 and 3.3 Å distance from the site was present and if they did not trigger and close contacts with protein or other solvent atoms. Following these modifications, refinement was repeated in *PHENIX*, and this procedure was iteratively performed until no more modifications to the model were warranted. In the latter cycles of refinement, individual B-factors, rather than group B-factors, were refined only if doing so lowered the free R-factor and narrowed the gap between R_work_ and R_free_. For papain-ligand complexes, as much refinement as was possible was performed prior to modeling any ligand in the active site. If evidence of a ligand was present in the F_o_-F_c_ map in the form of strong peaks stemming from the active cysteine 25 residue and exceeding 3 sigma levels, the ligand was modeled prior to the final cycles of refinement. A 1.8 Å restraint, with an allowed standard deviation of 0.02 Å, on the bond distance between the sulfur atom of Cys25 and the covalently bound carbon atom of the ligand was applied in *PHENIX*, and the occupancies of all ligand atoms were refined together as a group.

### Native mass spectrometry of on-grid material

On-grid material was harvested by removing the grid from liquid nitrogen, allowing it to thaw, and immediately pipetting 7µL of 150mM ammonium acetate onto the grid surface. Repeated pipetting was done to dislodge and dissolve material on the grid, and once the sample was fully dissolved it was subjected to nMS analysis. Samples were measured using a Thermo Q Exactive Plus UHMR Orbitrap instrument (Thermo Fisher Scientific, Bremen, Germany). NanoESI glass capillaries were pulled and coated with gold in-house. Samples were loaded into the gold coated capillary tubes and a 1.0-1.4 kV voltage was applied to the nanoESI source. All mass spectra are collected at a resolution of 12,500 at *m/z* 400 and instrument tuning parameters are listed in Table S14. Native mass spectra were deconvoluted using UniDec (version 7.0.2), a software tool for Bayesian deconvolution of mass and ion mobility spectra (Marty *et al*., 2015). Spectra were imported as thermo raw data and processed with the following parameters: a mass range of 5000 - 50000 Da, a charge state range of 1-20, an *m/z* range from 2000 - 4000, and masses were sampled every 1 Da.

### Single crystal X-ray diffraction and nMS of macroscopic crystals

Crystals of papain, both unliganded and in complex with ligands, were mounted on Mitegen loops and flash-frozen under a nitrogen stream at 100 K. Complete X-ray diffraction datasets were collected using a Rigaku FRE+ rotating anode X-ray diffractometer equipped with a Cu Kα source (λ = 1.54 Å) and a Rigaku HTC detector. Data were acquired with 2-minute integrated exposures over 0.5° oscillations per frame, with total collection times of approximately 26 hours per crystal. The detector was positioned at a distance of 78 mm from the crystal, mapping a resolution of 1.4 Å at the edge of the detector field.

Equivalent crystals were prepared for nMS by sequentially washing them in three 10 µL aliquots of 500 mM ammonium acetate to remove excess crystallization solution. Following the washes, crystals were dissolved in 10 µL of a working buffer containing 150 mM ammonium acetate to promote complete dissolution. The resulting samples were centrifuged at 10,000 × g for 5 minutes to ensure the removal of any residual crystal fragments. The supernatant was then transferred into glass capillaries for analysis.

### Fixed-target serial synchrotron X-ray crystallography data collection and processing

All SSX experiments were performed at the European Synchrotron Radiation Facility (ESRF) at beamline ID29 (Orlans *et al*., 2025). 5 µL of a concentrated slurry of papain microcrystals, either in apo-form or pre-soaked with a solution of E-64 or natural product analogs, was loaded between two faces of mylar foil and allowed to spread by capillary action. This film was loaded onto a fixed target chip and rastered while illuminated by a 11.56 keV X-ray beam delivering a series of 90 µs pulses at > 1.2 x 10^15^ photons per second flux. Diffraction was by the *MXCuBE-Web* software (Oscarsson *et al*., 2019) on a Jungfrau 4M detector, and frames containing diffraction were identified in real-time by the *LimA2* software (Debionne *et al*., 2023). Reflections from these hits were indexed using the *MOSFLM* (Battye *et al*., 2011) and *xGandalf* routines (Gevorkov *et al*., 2019) facilitated by *CrystFEL* (White *et al*., 2016), with the most likely known unit cell and space group symmetry for papain enforced. Data was scaled in *CrystFEL*, after which phasing and refinement were performed by the same protocol as described above for single-crystal data.

### LCMS of E-64 and natural product analogs

E-64 and analogs were diluted to a final concentration of 1 mg/ml, and 1uL injections were used for each run. Samples were separated by reversed-phase liquid chromatography through a C18 column (Poroshell 120 HPLC Column 2.7 μm 3.0 x 50mm) with a gradient flow from 0% ACN 0.1% formic acid to 100% ACN 0.1% formic acid at a flow rate of 0.6ml/min over a course of 15 minutes. The samples were analyzed with an Agilent 6545 Q-TOF LC/MS in positive ion mode with 1260 Infinity LC.

### Intact protein LCMS

Samples were crystallized, and papain was purified using the same method employed for EDMS. For the preparation, samples were incubated at 57°C for 1 hour under either reducing conditions (2 mM TCEP) or non-reducing conditions (no TCEP). Subsequently, 4 mM iodoacetamide was added to alkylate the protein, and the reaction was carried out for 30 minutes in the dark. The reaction was quenched with 8 mM TCEP. The final reaction volume was adjusted to 30 µL, with the protein concentration standardized to 1 mg/mL as determined by NanoDrop absorbance at A280. For LC-MS analysis, 5 µL of the 1 mg/mL sample was injected onto a C18 column (ZORBAX Reversed-Phase 2.7 μm 2.1 x 150mm). Separation was performed using a gradient elution from 0% ACN 0.1% formic acid to 100% ACN 0.1% formic acid at a flow rate of 0.8ml/min over 15 minutes on an Agilent 6530 Q-TOF LC/MS system in positive ion mode coupled with a 1260 Infinity LC system.

### Papain-inhibitor activity inhibition assay

To test the inhibition efficiency of natural product E-64 analogs against papain, a series of cleavage inhibition assays were performed against the 150 kDa antibody OKT9, is cleaved by papain. Solutions of OKT9, twice-crystallized papain extract, and E-64 analogs were diluted to working concentrations of 1 mg/mL, 1.25 mg/mL, and 2 mM respectively in digestion buffer (20 mM sodium phosphate pH 7.0, 10 mM EDTA). Reactions were carried out at a 1:10 papain:OTK9 mass ratio in 25 μL volumes testing cleavage ability in the either the absence of E-64 analogs, or the presence of E-64 analogs at 0.4 mM concentration. Reactions were carried out at 37 °C for 5-6 hours before quenching by snap freezing the mixture in liquid nitrogen. Cleavage activity was assessed by SDS-PAGE, comparing inhibited and uninhibited lanes to OKT9 untreated with papain, where disappearance of a band at the 150 kDa position indicating OKT9 was taken as evidence of papain activity.

### Synthesis of crude E-64 and E-64 analogs

Biosynthetic ligands were generated in one-pot reactions by adding precursor compounds to a solution containing the bioactive enzymes Cp1B and Cp1D, which catalyze the 2-step reaction resulting in E-64 and its analogs. For the production of E-64, 100 uL reactions containing 50 mM sodium phosphate buffer 8.0, 25 μM Cp1B, 25 μM Cp1D, 5 mM (±)-*trans*-epoxy-succinic acid, 2.5 μM L-leucine, 5mM agmatine, 10 mM ATP, and 10 mM MgCl2 were incubated at 30 oC for 16 hours. Reactions producing E-64-A65 were prepared identically to those for E-64, but substituting the L-leucine group with 2.5mM 2-amino-2-cyclobutylacetic acid and agmatine with 5 mM 1,8-diaminooctane. Following reaction completion, crude mixtures were spun through a 30 kDa Amicon concentrator tube to filter out the Cp1B and Cp1D enzymes. The flowthrough from these spins containing the product inhibitors were used in subsequent experiments.

### ArrayED ligand printing and on-grid soaking

Using a Scienion S3 microarrayer, droplets of a ~26 mM solution of E-64 (250−350 pL volume per drop) were automatically printed onto Quantifoil R2/2 Formvar/Carbon 200 Mesh Cu TEM grids in a 12×12 pattern, with one droplet per grid square (Fig. S7A,B). After drying, the printing process was repeated on the same grid to increase the amount of E-64 present. The grids were allowed to dry and stored under ambient conditions at room temperature until used for mixing with papain crystals and freezing. 2 μL of 66% methanol was added to the arrayed face of the grid and mixed for approximately 1 minute by repeated pipetting. Grids were mounted in an FEI Vitrobot with a humidity-saturated chamber held at12oC. 1 μL of 66% methanol was loaded on the back side of each grid, and 1.5 μL crystal slurry pre-treated with DTT was loaded on the front. The microcrystal slurry was allowed to mix with E-64 by incubating on the grid within the Vitrobot chamber for 5 minutes. After this waiting period, the grid was manually back blotted as described above, and plunge frozen into liquid ethane (Fig. S7A).

## Data Availability

MicroED structures of papain and papain-ligand complexes are available via PDB entries: 9NAG, 9NAE, 9NAR, 9NAO, 9N9D, 9NBQ, 9NC1, 9NAX, 9NAY, and 9NBP. MicroED structure of E-64D is available via CCDC deposition number 2423833. Single crystal X-ray diffraction structures of papain-ligand complexes are available via PDB entries: 9NB2 and 9NAT. SSX structures of papain and papain-ligand complexes are available via PDB entries: 9NCA, 9NBF, 9NBJ, 9NBK, 9NBN, 9NB4, and 9NB7. Diffraction data are available via Zenodo entries: 14876608 (MicroED data) and 14876818 (single crystal X-ray diffraction data). SSX data is available on the ESRF data portal under DOI 10.15151/ESRF-DC-2045905466. Files associated with additional structures referenced in this report but not deposited in the PDB are available via Zenodo entry 14885217.

## Acknowledgements

We thank Dr. Duilio Cascio and Dr. David Delgadillo for technical assistance. We acknowledge the European Synchrotron Radiation Facility (ESRF) for provision of synchrotron radiation facilities under ID29 BAG proposal MX2577. J.A.R. is supported as a Packard Fellow. This work was supported by the Howard Hughes Medical Institute Emerging Pathogens Initiative (HHMI-EPI), by DOE Grant DE-FC02-02ER63421, and performed as part of STROBE, an NSF Science and Technology Center through Grant DMR-1548924. The work was also supported by National Institutes of Health grants NIH-NIGMS Grant R35 GM128867, R35 GM145286, the NIH Ruth L. Kirschstein National Research Service Award program (GM007185), and T32GM145388.

## Author Contributions

N.W.V. designed experiments, prepared samples, collected and analyzed diffraction data; C.W.F., J.A.S., M.R.S., J.A.R. and S.W. collected and/or analyzed diffraction data; C.W.F. collected and analyzed mass spectrometry data with input from Y.C.; M.L., M.P.A., C.V.K., N.A. and P.R. prepared and characterized biosynthetic samples; S.J. printed arrayed ligand TEM samples; S.R., J.O., S.B., and D.D. assisted with SSX data collection; J.A.R. directed the work with input from Y.T., H.M.N, S.W. and J.A.L.; N.W.V. and J.A.R. wrote the manuscript with input from all authors.

