## Supplementary Information for "Combining MicroED and native mass spectrometry for structural discovery of enzyme-biosynthetic inhibitor complexes"

**This PDF file includes:**

Figures S1 to S17

Tables S1 to S13

CheckCIF structure validation report and response

**Other supporting materials for this manuscript include the following:**

Datasets deposited on *Zenodo*

Structures deposited to the PDB and CCDC

PDB structure validation reports

### Supplementary Tables

**Table S1.** Statistics of crystallographic data reduction and refinement for MicroED structures of apo-form papain, papain co-crystallized with E-64, and papain co-crystallized with E-64-A65.

| | Papain (apo) | Papain-E-64 | Papain-E-64-A65 |
|---|---|---|---|
| **PDB ID** | 9NAG | 9NAE | 9NAO |
| **Data Collection and Processing** | | | |
| No. crystals merged | 2 | 1 | 1 |
| Temperature (K) | 100 | 100 | 100 |
| Electron wavelength (Å) | 0.0251 | 0.0251 | 0.0251 |
| Resolution (Å) | 50.08 – 2.50 (2.60-2.50) | 50.34- 2.30 (2.40-2.30) | 50.24 – 2.50 (2.60 – 2.50) |
| Space Group | $P2_12_12_1$ | $P2_12_12_1$ | $P2_12_12_1$ |
| *a*, *b*, *c* (Å) | 41.99, 49.09, 100.15 | 42.18, 49.25, 100.69 | 42.31, 48.32, 100.48 |
| *α*, *β*, *γ* (°) | 90,90,90 | 90,90,90 | 90,90,90 |
| # total reflections | 33883 (3776) | 30859 (3725) | 24784 (2817) |
| # unique reflections | 6918 (759) | 9018 (1062) | 6665 (742) |
| $R_{merge}$ (%) | 29.5 (47.4) | 25.3 (103.6) | 24.9 (103.4) |
| CC1/2 (%) | 94.6 (42.8) | 96.6 (41.2) | 96.2 (45.5) |
| <I/σI> | 5.48 (3.62) | 3.84 (1.53) | 4.27 (1.36) |
| Completeness (%) | 90.5 (91.1) | 91.4 (92.6) | 87.7 (91.2) |
| **Phasing** | | | |
| Search model PDB | 9PAP | 9PAP | 9PAP |
| **Refinement** | | | |
| Resolution (Å) | 50.08 – 2.80 | 50.34 – 2.30 | 50.24 – 2.50 |
| $R_{work}$ (%) | 21.28 | 20.31 | 23.35 |
| $R_{free}$ (%) | 24.78 | 25.70 | 26.46 |
| # protein atoms | 1720 | 1699 | 1717 |
| # ligand atoms | 0 | 25 | 26 |
| # solvent atoms | 13 | 22 | 9 |

| | | | |
|---|---|---|---|
| Occupancy of ligand atoms (%) | N/A | 73 | 67 |
| Average B-factor (Å$^2$) | | | |
| Protein | 28.34 | 23.84 | 31.14 |
| Ligand | N/A | 30.62 | 37.44 |
| Solvent | 10.95 | 19.17 | 18.09 |
| Ramachandran (%) | | | |
| Favored (%) | 97.62 | 98.10 | 97.14 |
| Allowed (%) | 2.38 | 1.90 | 2.86 |
| Outliers (%) | 0.00 | 0.00 | 0.00 |
| R.M.S. deviations | | | |
| Bond lengths (Å) | 0.007 | 0.007 | 0.007 |
| Bond angles (°) | 0.999 | 0.959 | 1.044 |
| Clashscore | 5.35 | 4.87 | 8.55 |

**Table S2.** Statistics of crystallographic data reduction and refinement for MicroED structures of papain soaked with E-64 for 30 seconds, 4 minutes, and 10 minutes. Structures marked with * did not feature adequately clear density to model a ligand, so the refinement statistics reported reference the model without a ligand modeled. The structure corresponding to the longest soaking time was deposited in the PDB.

| | Papain-E-64 (30 seconds)* | Papain-E-64 (4 minutes) | Papain-E-64 (10 minutes) |
|---|---|---|---|
| **PDB ID** | N/A | N/A | 9NAR |
| **Data Collection and Processing** | | | |
| No. crystals merged | 1 | 2 | 2 |
| Temperature (K) | 100 | 100 | 100 |
| Electron wavelength (Å) | 0.0251 | 0.0251 | 0.0251 |
| Resolution (Å) | 26.85 – 2.50 (2.60 -2.50) | 49.98 – 2.20 (2.30 – 2.20) | 50.12 – 2.50 (2.60 – 2.50) |
| Space Group | $P2_12_12_1$ | $P2_12_12_1$ | $P2_12_12_1$ |
| *a*, *b*, *c* (Å) | 41.28, 49.36, 101.28 | 42.00, 49.39, 99.95 | 41.92, 48.32, 100.24 |
| *α*, *β*, *γ* (°) | 90,90,90 | 90,90,90 | 90,90,90 |
| # total reflections | 41792 (4814) | 70608 (8476) | 41982 (4636) |
| # unique reflections | 6708 (732) | 11081 (1335) | 7342 (790) |
| $R_{merge}$ (%) | 33.2 (56.0) | 31.3 (83.6) | 29.6 (112.8) |
| CC1/2 (%) | 96.7 (29.0) | 97.7 (55.6) | 97.1 (55.4) |
| $\langle I/\sigma I \rangle$ | 5.44 (3.41) | 5.76 (2.49) | 6.31 (2.19) |
| Completeness (%) | 87.9 (88.5) | 99.4 (97.9) | 97.7 (98.5) |
| **Phasing** | | | |
| Search model PDB | 9PAP | 9PAP | 9PAP |
| **Refinement** | | | |
| Resolution (Å) | 26.85 – 2.50 | 49.98 – 2.50 | 50.12 – 2.50 |
| $R_{work}$ (%) | 23.09 | 18.76 | 19.24 |
| $R_{free}$ (%) | 27.10 | 24.64 | 20.59 |
| # protein atoms | 1716 | 1711 | 1680 |
| # ligand atoms | 0 | 25 | 25 |

| | | | |
|---|---|---|---|
| # solvent atoms | 6 | 41 | 31 |
| Occupancy of ligand atoms (%) | N/A | 72 | 77 |
| Average B-factor (Å$^2$) | | | |
| Protein | 14.57 | 18.77 | 21.80 |
| Ligand | N/A | 29.45 | 23.98 |
| Solvent | 3.38 | 12.42 | 12.74 |
| Ramachandran (%) | | | |
| Favored (%) | 97.14 | 98.10 | 97.62 |
| Allowed (%) | 2.86 | 1.90 | 2.38 |
| Outliers (%) | 0.00 | 0.00 | 0.00 |
| R.M.S. deviations | | | |
| Bond lengths (Å) | 0.011 | 0.008 | 0.007 |
| Bond angles (°) | 1.382 | 0.971 | 0.935 |
| Clashscore | 11 | 6.18 | 5.08 |

**Table S3.** Statistics of crystallographic data reduction and refinement for MicroED structures of papain soaked with ligand mixtures. For structures marked with *, active-site ligand density was not sufficiently clear for confident ligand modeling, so ligands were not included in refinement for these structures and they were not deposited in the PDB.

| | Papain-protease inhibitor cocktail | Papain-E-64,E-64C, and E-64D mixture | Papain-E-64-A65, E315, E372, and E405 mixture* | Papain-E315, E372, and E405 mixture* |
|---|---|---|---|---|
| **PDB ID** | 9NC1 | 9NCA | N/A | N/A |
| **Data Collection and Processing** | | | | |
| No. crystals merged | 2 | 3 | 1 | 1 |
| Temperature (K) | 100 | 100 | 100 | 100 |
| Electron wavelength (Å) | 0.0251 | 0.0251 | 0.0251 | 0.0251 |
| Resolution (Å) | 32.06 – 2.40 (2.50 – 2.40) | 49.82 – 2.50 (2.60 – 2.50) | 19.86 – 2.40 (2.50 – 2.40) | 31.86 – 2.50 (2.60 – 2.50) |
| Space Group | $P2_12_12_1$ | $P2_12_12_1$ | $P2_12_12_1$ | $P2_12_12_1$ |
| *a*, *b*, *c* (Å) | 42.57, 48.85, 99.86 | 42.74, 48.96, 99.64 | 42.22, 49.15, 101.25 | 41.83, 49.18, 99.99 |
| $\alpha$, $\beta$, $\gamma$ (°) | 90,90,90 | 90,90,90 | 90,90,90 | 90,90,90 |
| # total reflections | 36267 (4240) | 49787 (5215) | 34102 (4020) | 26915 (3040) |
| # unique reflections | 8231 (940) | 6282 (638) | 7088 (806) | 6628 (733) |
| $R_{merge}$ (%) | 24.6 (67.2) | 33.5 (120.7) | 38.2 (126.0) | 26.4 (95.6) |
| CC1/2 (%) | 96.7 (58.8) | 86.7 (62.3) | 95.9 (42.0) | 95.7 (35.5) |
| $\langle I/\sigma I \rangle$ | 5.31 (2.63) | 6.12 (2.45) | 4.69 (1.90) | 5.03 (1.79) |
| Completeness (%) | 95.2 (96.7) | 81.5 (76.9) | 81.3 (81.3) | 87.0 (87.8) |
| **Phasing** | | | | |
| Search model PDB | 9PAP | 9PAP | 9PAP | 9PAP |
| **Refinement** | | | | |
| Resolution (Å) | 32.06 – 2.40 | 49.82 – 2.50 | 19.86 – 2.40 | 31.86 – 2.50 |
| $R_{work}$ (%) | 20.33 | 22.16 | 23.65 | 20.75 |
| $R_{free}$ (%) | 26.09 | 27.56 | 28.27 | 26.07 |
| # protein atoms | 1655 | 1681 | 1674 | 1695 |
| # ligand atoms | 12 | 47 | 0 | 0 |

| | | | | |
|---|---|---|---|---|
| # solvent atoms | 25 | 2 | 13 | 13 |
| Occupancy of ligand atoms (%) | 76 | 52 (E-64), 42 (E-64C) | N/A | N/A |
| Average B-factor (Å$^2$) | | | | |
| Protein | 22.83 | 18.10 | 27.65 | 25.46 |
| Ligand | 29.16 | 38.04 | N/A | N/A |
| Solvent | 14.70 | 2.83 | 24.10 | 13.22 |
| Ramachandran (%) | | | | |
| Favored (%) | 97.62 | 98.10 | 97.14 | 96.67 |
| Allowed (%) | 2.38 | 1.90 | 2.86 | 3.33 |
| Outliers (%) | 0.00 | 0.00 | 0.00 | 0.00 |
| R.M.S. deviations | | | | |
| Bond lengths (Å) | 0.006 | 0.009 | 0.013 | 0.011 |
| Bond angles (°) | 0.914 | 1.311 | 1.606 | 1.472 |
| Clashscore | 2.43 | 10.68 | 8.55 | 9.33 |

**Table S4.** Statistics of crystallographic data reduction and refinement for MicroED structures of papain soaked with crude preparations of E-64 and E-64-A65

| | Papain-E-64 (crude) | Papain-E-64-A65 (crude) |
|---|---|---|
| **PDB ID** | 9NAX | 9NAY |
| **Data Collection and Processing** | | |
| No. crystals merged | 2 | 2 |
| Temperature (K) | 100 | 100 |
| Electron wavelength (Å) | 0.0251 | 0.0251 |
| Resolution (Å) | 49.93 – 2.30 (2.40 – 2.30) | 50.05 – 2.50 (2.60 – 2.50) |
| Space Group | $P2_12_12_1$ | $P2_12_12_1$ |
| *a*, *b*, *c* (Å) | 41.85, 48.63, 99.87 | 42.20, 48.78, 100.10 |
| *α*, *β*, *γ* (°) | 90,90,90 | 90,90,90 |
| # total reflections | 69072 (7944) | 40867 (4205) |
| # unique reflections | 8337 (958) | 6582 (688) |
| $R_{merge}$ (%) | 28.0 (88.2) | 27.5 (93.7) |
| CC1/2 (%) | 98.1 (66.3) | 97.0 (55.4) |
| <I/σI> | 6.46 (2.55) | 5.46 (1.92) |
| Completeness (%) | 87.0 (87.6) | 86.4 (85.4) |
| **Phasing** | | |
| Search model PDB | 9PAP | 9PAP |
| **Refinement** | | |
| Resolution (Å) | 49.93 – 2.30 | 50.05 – 2.50 |
| $R_{work}$ (%) | 18.99 | 17.97 |
| $R_{free}$ (%) | 24.75 | 22.77 |
| # protein atoms | 1666 | 1689 |
| # ligand atoms | 25 | 26 |
| # solvent atoms | 25 | 4 |

| | | |
|---|---|---|
| Occupancy of ligand atoms (%) | 75 | 75 |
| Average B-factor (Å$^2$) | | |
| Protein | 25.08 | 29.41 |
| Ligand | 31.38 | 41.61 |
| Solvent | 18.05 | 13.83 |
| Ramachandran (%) | | |
| Favored (%) | 97.62 | 98.10 |
| Allowed (%) | 2.38 | 1.90 |
| Outliers (%) | 0.00 | 0.00 |
| R.M.S. deviations | | |
| Bond lengths (Å) | 0.007 | 0.007 |
| Bond angles (°) | 0.964 | 1.119 |
| Clashscore | 5.13 | 7.50 |

**Table S5.** Statistics of crystallographic data reduction and refinement for MicroED structures of papain co-crystallized with E-64C and E-64D.

| | Papain-E-64C | Papain-E-64D |
|---|---|---|
| **PDB ID** | 9N9D | 9NBQ |
| **Data Collection and Processing** | | |
| No. crystals merged | 1 | 1 |
| Temperature (K) | 100 | 100 |
| Electron wavelength (Å) | 0.0251 | 0.0251 |
| Resolution (Å) | 50.46 – 2.20 (2.30 – 2.20) | 32.32 – 2.30 (2.40 – 2.30) |
| Space Group | $P2_12_12_1$ | $P2_12_12_1$ |
| *a*, *b*, *c* (Å) | 41.88, 48.93, 100.93 | 42.08, 49.34, 100.92 |
| *α*, *β*, *γ* (°) | 90,90,90 | 90,90,90 |
| # total reflections | 43354 (5189) | 37992 (4444) |
| # unique reflections | 10957 (1314) | 9338 (1076) |
| $R_{merge}$ (%) | 26.8 (98.8) | 34.4 (94.6) |
| CC1/2 (%) | 97.0 (45.9) | 94.3 (47.4) |
| <I/σI> | 5.29 (2.00) | 4.68 (2.04) |
| Completeness (%) | 98.7 (97.8) | 94.5 (93.6) |
| **Phasing** | | |
| Search model PDB | 9PAP | 9PAP |
| **Refinement** | | |
| Resolution (Å) | 50.46 – 2.20 | 32.32 – 2.30 |
| $R_{work}$ (%) | 18.12 | 19.33 |
| $R_{free}$ (%) | 23.63 | 24.70 |
| # protein atoms | 1729 | 1724 |
| # ligand atoms | 41 | 24 |
| # solvent atoms | 23.25 | 33 |
| Occupancy of ligand atoms (%) | 81 | 74 |

| | | |
|---|---|---|
| Average B-factor ($Å^2$) | | |
| Protein | 23.00 | 19.29 |
| Ligand | 39.02 | 43.79 |
| Solvent | 17.46 | 12.32 |
| Ramachandran (%) | | |
| Favored (%) | 97.14 | 97.62 |
| Allowed (%) | 2.86 | 2.38 |
| Outliers (%) | 0.00 | 0.00 |
| R.M.S. deviations | | |
| Bond lengths (Å) | 0.009 | 0.007 |
| Bond angles (°) | 1.327 | 0.958 |
| Clashscore | 4.36 | 6.41 |

**Table S6.** Statistics of crystallographic data reduction and refinement for MicroED structure of papain from microcrystals mixed on-grid with arrayed E-64.

| | Papain-E-64 (array sample preparation) |
|---|---|
| **PDB ID** | 9NBP |
| **Data Collection and Processing** | |
| No. crystals merged | 2 |
| Temperature (K) | 100 |
| Electron wavelength (Å) | 0.0251 |
| Resolution (Å) | – 2.80 (2.90 – 2.80) |
| Space Group | $P2_12_12_1$ |
| *a*, *b*, *c* (Å) | 41.70, 49.28, 100.44 |
| *α*, *β*, *γ* (°) | 90,90,90 |
| # total reflections | 25374 (2426) |
| # unique reflections | 4950 (471) |
| $R_{merge}$ (%) | 37.6 (95.8) |
| CC1/2 (%) | 94.8 (62.3) |
| <I/σI> | 4.74 (2.02) |
| Completeness (%) | 90.1 (89.9) |
| **Phasing** | |
| Search model PDB | 9PAP |
| **Refinement** | |
| Resolution (Å) | – 2.80 |
| $R_{work}$ (%) | 19.79 |
| $R_{free}$ (%) | 24.61 |
| # protein atoms | 1682 |
| # ligand atoms | 10 |
| # solvent atoms | 25 |
| Occupancy of ligand atoms (%) | 75 |
| Average B-factor (Å$^2$) | |
| Protein | 19.82 |
| Ligand | 26.53 |

| | |
|---|---|
| Solvent | 7.15 |
| Ramachandran (%) | |
| Favored (%) | 97.62 |
| Allowed (%) | 2.38 |
| Outliers (%) | 0.00 |
| R.M.S. deviations | |
| Bond lengths (Å) | 0.007 |
| Bond angles (°) | 1.014 |
| Clashscore | 7.47 |

**Table S7.** Statistics of crystallographic data reduction and refinement for single-crystal XRD structures of papain soaked with E-64 for 30 seconds, 4 minutes, 10 minutes, 80 minutes, and 72 hours. Structures marked with * did not feature adequately clear density to model a ligand, so the refinement statistics reported reference the model without a ligand modeled. The structure corresponding to the longest soaking time was deposited in the PDB.

| | Papain-E-64 (30 seconds)* | Papain-E-64 (4 minutes)* | Papain-E-64 (10 minutes)* | Papain-E-64 (80 minutes) | Papain-E-64 (72 hours) |
|---|---|---|---|---|---|
| **PDB ID** | N/A | N/A | N/A | N/A | 9NB2 |
| **Data Collection and Processing** | | | | | |
| X-ray wavelength (Å) | 1.54 | 1.54 | 1.54 | 1.54 | 1.54 |
| Temperature (K) | 100 | 100 | 100 | 100 | 100 |
| Resolution (Å) | 50.80 – 1.50 (1.60 – 1.50) | 50.81 – 1.50 (1.60 – 1.50) | 39.21 – 1.50 (1.60 – 1.50) | 44.23 – 1.60 (1.70 – 1.60) | 27.86 – 1.50 (1.60 – 1.50) |
| Space Group | $P2_12_12_1$ | $P2_12_12_1$ | $P2_12_12_1$ | $P2_12_12_1$ | $P2_12_12_1$ |
| *a*, *b*, *c* (Å) | 42.65, 49.18, 101.61 | 42.57, 49.10, 101.62 | 42.51, 49.27, 101.54 | 42.57, 49.15 101.38 | 42.41, 48.91, 101.71 |
| *α*, *β*, *γ* (°) | 90,90,90 | 90,90,90 | 90,90,90 | 90,90,90 | 90,90,90 |
| # total reflections | 440990 (77330) | 442445 (76714) | 458274 (78829) | 355576 (63574) | 417970 (71570) |
| # unique reflections | 34702 (5895) | 34861 (6005) | 34970 (6033) | 27776 (4456) | 34540 (6000) |
| $R_{merge}$ (%) | 11.3 (99.2) | 15.1 (83.3) | 7.2 (55.0) | 5.6 (36.4) | 7.8 (71.0) |
| CC1/2 (%) | 99.8 (89.5) | 99.6 (67.0) | 99.9 (92.1) | 99.9 (96.7) | 99.9 (85.0) |
| $\langle I/\sigma I \rangle$ | 12.53 (2.95) | 11.29 (3.37) | 20.53 (4.93) | 26.44 (6.13) | 18.22 (3.72) |
| Completeness (%) | 98.8 (97.0) | 99.8 (99.3) | 99.8 (99.6) | 96.1 (94.1) | 99.4 (100.0) |
| **Phasing** | | | | | |
| Search model PDB | 9PAP | 9PAP | 9PAP | 9PAP | 9PAP |
| **Refinement** | | | | | |
| Resolution (Å) | 50.80 – 1.50 | 50.80 – 1.50 | 39.21 – 1.50 | 44.23 – 1.60 | 27.86 – 1.50 |
| $R_{work}$ (%) | 19.71 | 19.33 | 18.95 | 22.00 | 18.35 |
| $R_{free}$ (%) | 22.30 | 21.53 | 20.75 | 26.51 | 21.24 |
| # protein atoms | 1721 | 1703 | 1706 | 1699 | 1725 |
| # ligand atoms | 0 | 0 | 0 | 25 | 25 |

| | | | | | |
|---|---|---|---|---|---|
| # solvent atoms | 213 | 242 | 283 | 211 | 230 |
| Occupancy of ligand atoms (%) | N/A | N/A | N/A | 61 | 82 |
| Average B-factor (Å$^2$) | | | | | |
| Protein | 20.34 | 17.36 | 18.29 | 19.63 | 18.41 |
| Ligand | N/A | N/A | N/A | 24.14 | 28.96 |
| Solvent | 29.18 | 26.48 | 29.71 | 27.38 | 26.95 |
| Ramachandran (%) | | | | | |
| Favored (%) | 98.07 | 98.55 | 98.07 | 98.10 | 98.57 |
| Allowed (%) | 1.93 | 1.45 | 1.93 | 1.90 | 1.43 |
| Outliers (%) | 0.00 | 0.00 | 0.00 | 0.00 | 0.00 |
| R.M.S. deviations | | | | | |
| Bond lengths (Å) | 0.006 | 0.006 | 0.006 | 0.007 | 0.006 |
| Bond angles (°) | 0.898 | 0.892 | 0.857 | 0.898 | 0.838 |
| Clashscore | 1.49 | 1.80 | 2.10 | 1.48 | 2.33 |

**Table S8.** Statistics of crystallographic data reduction and refinement for single-crystal XRD structure of papain soaked with a commercial protease inhibitor cocktail for 20 hours. Refinement statistics reported are for the protein without ligand modeled, as the ligand density could not be unambiguously assigned.

| | Papain-protease inhibitor cocktail (XRD) |
|---|---|
| **PDB ID** | N/A |
| **Data Collection and Processing** | |
| X-ray wavelength (Å) | 1.54 |
| Temperature (K) | 100 |
| Resolution (Å) | 44.03 – 1.50 (1.60 – 1.50) |
| Space Group | $P2_12_12_1$ |
| *a*, *b*, *c* (Å) | 42.37, 48.87, 101.49 |
| *α*, *β*, *γ* (°) | 90,90,90 |
| # total reflections | 459015 (78661) |
| # unique reflections | 34593 (5971) |
| $R_{merge}$ (%) | 11.8 (72.0) |
| CC1/2 (%) | 99.7 (89.7) |
| <I/σI> | 13.84 (3.42) |
| Completeness (%) | 100.9 (100.0) |
| **Phasing** | |
| Search model PDB | 9PAP |
| **Refinement** | |
| Resolution (Å) | 44.03 – 1.50 |
| $R_{work}$ (%) | 21.41 |
| $R_{free}$ (%) | 23.68 |
| # protein atoms | 1652 |
| # ligand atoms | 0 |
| # solvent atoms | 108 |
| Occupancy of ligand atoms (%) | N/A |

| | |
|---|---|
| Average B-factor (Å$^2$) | |
| Protein | 20.76 |
| Ligand | N/A |
| Solvent | 24.36 |
| Ramachandran (%) | |
| Favored (%) | 97.62 |
| Allowed (%) | 2.38 |
| Outliers (%) | 0.00 |
| R.M.S. deviations | |
| Bond lengths (Å) | 0.006 |
| Bond angles (º) | 1.047 |
| Clashscore | 2.47 |

**Table S9.** Statistics of crystallographic data reduction and refinement for single-crystal XRD structure of papain co-crystallized with leupeptin.

| | Papain-leupeptin (XRD) |
|---|---|
| **PDB ID** | 9NAT |
| **Data Collection and Processing** | |
| X-ray/electron wavelength (Å) | 1.54 |
| Temperature (K) | 100 |
| Resolution (Å) | 39.11 – 1.60 (1.70 – 1.60) |
| Space Group | $P2_12_12_1$ |
| *a*, *b*, *c* (Å) | 42.45, 48.91, 100.80 |
| *α*, *β*, *γ* (°) | 90,90,90 |
| # total reflections | 380919 (62121) |
| # unique reflections | 28509 (4665) |
| $R_{merge}$ (%) | 6.5 (16.9) |
| CC1/2 (%) | 99.9 (99.4) |
| <I/σI> | 26.69 (11.90) |
| Completeness (%) | 100 (100) |
| **Phasing** | |
| Search model PDB | 9PAP |
| **Refinement** | |
| Resolution (Å) | 39.11 – 1.60 |
| $R_{work}$ (%) | 16.92 |
| $R_{free}$ (%) | 19.76 |
| # protein atoms | 1708 |
| # ligand atoms | 30 |
| # solvent atoms | 267 |
| Occupancy of ligand atoms (%) | 75 |
| Average B-factor (Å$^2$) | |

| | |
|---|---|
| Protein | 15.44 |
| Ligand | 28.28 |
| Solvent | 24.87 |
| Ramachandran (%) | |
| Favored (%) | 98.10 |
| Allowed (%) | 1.90 |
| Outliers (%) | 0.00 |
| R.M.S. deviations | |
| Bond lengths (Å) | 0.006 |
| Bond angles (°) | 0.936 |
| Clashscore | 1.48 |

**Table S10.** Statistics of crystallographic data reduction and refinement for serial synchrotron X-ray diffraction structures of apo-form papain, papain soaked with E-64, papain soaked with E-64C, papain soaked with E-64D, and papain soaked with an equimolar mixture of all three ligands. For the mixture soaking experiment, the common structural fragment between the three potential ligands was modeled in the structure and refined.

| | Papain (apo) | Papain-E-64 | Papain-E-64C | Papain-E-64D | Papain-E-64,E-64C,E-64D mixture |
|---|---|---|---|---|---|
| **PDB ID** | 9NCC | 9NBF | 9NBJ | 9NBK | 9NBN |
| **Data Collection and Processing** | | | | | |
| X-ray wavelength (Å) | 1.072 | 1.072 | 1.072 | 1.072 | 1.073 |
| Temperature (K) | 293 | 293 | 293 | 293 | 293 |
| # hits | 9705 | 7857 | 22890 | 41210 | 23328 |
| # indexed images | 7284 | 6390 | 18301 | 33457 | 15005 |
| # crystals indexed | 8201 | 6921 | 19985 | 36673 | 16147 |
| Resolution (Å) | 44.05 – 1.80 (1.83 – 1.80) | 44.05 – 1.80 (1.83 – 1.80) | 44.05 – 1.80 (1.83 – 1.80) | 44.05 – 1.80 (1.83 – 1.80) | 44.05 – 1.80 (1.83 – 1.80) |
| Space Group | $P2_12_12_1$ | $P2_12_12_1$ | $P2_12_12_1$ | $P2_12_12_1$ | $P2_12_12_1$ |
| *a*, *b*, *c* (Å) | 42.5, 48.9, 101.5 | 42.5, 48.9, 101.5 | 42.5, 48.9, 101.5 | 42.5, 48.9, 101.5 | 42.5, 48.9, 101.5 |
| *α*, *β*, *γ* (°) | 90,90,90 | 90,90,90 | 90,90,90 | 90,90,90 | 90,90,90 |
| # total reflections | 3195456 (109871) | 2591698 (90854) | 9961279 (349481) | 18939476 (661124) | 4617351 (162646) |
| # unique reflections | 37921 (1877) | 37298 (1877) | 37928 (1877) | 37928 (1877) | 37928 (1877) |
| $R_{split}$ (%) | 31.7 (212.7) | 37.5 (239.5) | 21.3 (86.5) | 15.4 (58.9) | 30.4 (239.2) |
| CC*(%) | 96.7 (46.9) | 94.7 (39.3) | 98.2 (76.3) | 99.1 (86.4) | 97.3 (39.5) |
| <I/σI> | 2.6 (0.5) | 2.3 (0.5) | 4.3 (1.2) | 6.1 (1.8) | 2.6 (0.5) |
| Completeness (%) | 100 (100) | 100 (100) | 100 (100) | 100 (100) | 100 (100) |
| **Phasing** | | | | | |
| Search model PDB | 9PAP | 9PAP | 9PAP | 9PAP | 9PAP |
| **Refinement** | | | | | |
| Resolution (Å) | 44.05 – 1.80 | 44.05 – 1.80 | 44.05 – 1.80 | 44.05 – 1.80 | 44.05 – 1.80 |

| | | | | | |
|---|---|---|---|---|---|
| $R_{work}$ (%) | 19.75 | 20.23 | 17.00 | 15.89 | 19.40 |
| $R_{free}$ (%) | 23.25 | 23.90 | 21.00 | 19.75 | 23.77 |
| # protein atoms | 1712 | 1714 | 1698 | 1709 | 1683 |
| # ligand atoms | 0 | 25 | 22 | 24 | 17 |
| # solvent atoms | 59 | 87 | 102 | 106 | 77 |
| Occupancy of ligand atoms (%) | N/A | 60 | 58 | 65 | 73 |
| Average B-factor (Å$^2$) | | | | | |
| Protein | 32.43 | 33.30 | 32.22 | 32.11 | 35.34 |
| Ligand | N/A | 39.80 | 42.11 | 44.31 | 42.68 |
| Solvent | 34.20 | 37.18 | 38.09 | 39.52 | 37.66 |
| Ramachandran (%) | | | | | |
| Favored (%) | 97.14 | 97.14 | 97.14 | 97.14 | 97.62 |
| Allowed (%) | 2.86 | 2.86 | 2.86 | 2.86 | 2.38 |
| Outliers (%) | 0.00 | 0.00 | 0.00 | 0.00 | 0.00 |
| R.M.S. deviations | | | | | |
| Bond lengths (Å) | 0.008 | 0.007 | 0.007 | 0.008 | 0.008 |
| Bond angles (°) | 0.983 | 0.939 | 0.958 | 0.954 | 0.959 |
| Clashscore | 4.47 | 4.69 | 4.15 | 3.81 | 3.31 |

**Table S11.** Statistics of crystallographic data reduction and refinement for serial synchrotron X-ray diffraction structures of papain soaked with E-64-A65, papain soaked with E315, papain soaked with E371, papain soaked with E405, and papain soaked with an equimolar mixture of all four natural product compounds. For structures marked with *, active-site ligand density was not sufficiently clear for confident ligand modeling, so ligands were not included in refinement for these structures and they were not deposited in the PDB.

| | Papain-E-64-A65 | Papain-E315* | Papain-371* | Papain-E405 | Papain-E-64-A65,E315,E371,E405 mixture* |
|---|---|---|---|---|---|
| **PDB ID** | 9NB4 | N/A | N/A | 9NB7 | N/A |
| **Data Collection and Processing** | | | | | |
| X-ray wavelength (Å) | 1.072 | 1.072 | 1.072 | 1.072 | 1.072 |
| Temperature (K) | 293 | 293 | 293 | 293 | 293 |
| # hits | 25777 | 37779 | 28180 | 27585 | 24241 |
| # indexed images | 22420 | 31349 | 20318 | 21818 | 20831 |
| # crystals indexed | 24052 | 33986 | 21621 | 23401 | 22368 |
| Resolution (Å) | 44.05 – 1.80 (1.83 – 1.80) | 44.05 – 1.80 (1.83 – 1.80) | 44.05 – 1.80 (1.83 – 1.80) | 44.05 – 1.80 (1.83 – 1.80) | 44.05 – 1.80 (1.83 – 1.80) |
| Space Group | $P2_12_12_1$ | $P2_12_12_1$ | $P2_12_12_1$ | $P2_12_12_1$ | $P2_12_12_1$ |
| *a*, *b*, *c* (Å) | 42.5, 48.9, 101.5 | 42.5, 48.9, 101.5 | 42.5, 48.9, 101.5 | 42.5, 48.9, 101.5 | 42.5, 48.9, 101.5 |
| *α*, *β*, *γ* (°) | 90,90,90 | 90,90,90 | 90,90,90 | 90,90,90 | 90,90,90 |
| # total reflections | 12627156 (441126) | 16706172 (582261) | 10335316 (359513) | 11270147 (394517) | 8021599 (281167) |
| # unique reflections | 37928 (1877) | 37928 (1877) | 37928 (1877) | 37928 (1877) | 37928 (1877) |
| $R_{split}$ (%) | 18.6 (79.6) | 16.6 (70.9) | 19.2 (124.2) | 20.6 (92.1) | 23.3 (23.3) |
| CC* (%) | 98.7 (77.0) | 99.0 (80.6) | 98.8 (64.4) | 98.3 (75.4) | 98.0 (74.6) |
| <I/σI> | 4.9 (1.3) | 5.6 (1.5) | 4.2 (0.9) | 4.6 (1.2) | 3.9 (1.1) |
| Completeness (%) | 100 (100) | 100 (100) | 100 (100) | 100 (100 | 100 (100) |
| **Phasing** | | | | | |
| Search model PDB | 9PAP | 9PAP | 9PAP | 9PAP | 9PAP |

**Refinement**

| | | | | | |
|---|---|---|---|---|---|
| Resolution (Å) | 44.05 – 1.80 | 44.05 – 1.80 | 44.05 – 1.80 | 44.05 – 1.80 | 44.05 – 1.80 |
| $R_{work}$ (%) | 16.67 | 16.29 | 17.48 | 17.39 | 19.33 |
| $R_{free}$ (%) | 20.96 | 19.68 | 20.23 | 21.50 | 23.28 |
| # protein atoms | 1721 | 1700 | 1669 | 1706 | 1652 |
| # ligand atoms | 26 | 0 | 0 | 29 | 0 |
| # solvent atoms | 95 | 83 | 102 | 79 | 49 |
| Occupancy of ligand atoms (%) | 68 | N/A | N/A | 64 | N/A |
| Average B-factor (Å$^2$) | | | | | |
| Protein | 31.79 | 32.13 | 32.82 | 31.93 | 31.76 |
| Ligand | 41.74 | N/A | N/A | 43.33 | N/A |
| Solvent | 36.25 | 35.82 | 38.96 | 37.22 | 31.13 |
| Ramachandran (%) | | | | | |
| Favored (%) | 98.57 | 97.14 | 96.67 | 97.62 | 97.14 |
| Allowed (%) | 1.43 | 2.86 | 3.33 | 2.38 | 2.86 |
| Outliers (%) | 0.00 | 0.00 | 0.00 | 0.00 | 0.00 |
| R.M.S. deviations | | | | | |
| Bond lengths (Å) | 0.007 | 0.008 | 0.008 | 0.007 | 0.007 |
| Bond angles (°) | 0.965 | 0.936 | 0.985 | 1.041 | 0.881 |
| Clashscore | 1.77 | 5.06 | 5.50 | 2.37 | 4.01 |

**Table S12.** Statistics of crystallographic data reduction and refinement for MicroED structure of E-64D

| | E-64D |
|---|---|
| **CCDC number** | 2423833 |
| **Data Collection and Processing** | |
| No. crystals merged | 1 |
| Electron wavelength (Å) | 0.0251 |
| Temperature (K) | 100 |
| Resolution (Å) | 12.08 – 0.8 (0.9 – 0.8) |
| Space Group | $P2_12_12_1$ |
| *a*, *b*, *c* (Å) | 4.72, 13.31, 28.78 |
| *α*, *β*, *γ* (°) | 90,90,90 |
| # total reflections | 8127 (2218) |
| # unique reflections | 1992 (551) |
| $R_{merge}$ (%) | 11.7 (34.9) |
| CC1/2 (%) | 99.3 (76.9) |
| <I/σI> | 7.02 (2.85) |
| Completeness (%) | 90.0 (89.4) |
| **Phasing** | |
| Software | *SHELXT* |
| **Refinement** | |
| Resolution (Å) | 12.08 – 0.8 |
| $R_1$ (%) | 15.75 |
| $wR_2$ (%) | 40.95 |
| Goodness of fit | 1.400 |

**Table S13.** Expected molecular weights of papain and ligands referenced for nMS analysis

| Species | Average Mass(Da) | Papain(reduced C25) + Ligand (Da) |
|---|---|---|
| Papain(Reduced C25) | 23422.9 | N/A |
| Papain(Sulfinic C25) | 23454.89 | N/A |
| E-64 | 357.2 | 23780.1 |
| E-64-A65 | 369.23 | 23792.13 |
| E-64C | 314.38 | 23737.28 |
| E-64D | 342.44 | 23765.34 |
| Leupeptin | 426.6 | 23849.5 |
| E315 | 315.18 | 23738.08 |
| E371 | 371.21 | 23794.11 |
| E405 | 405.19 | 23828.09 |

### Table S14 Instrument tuning parameters for ion transmission

| | |
|---|---|
| **Instrument parameters** | |
| Ion mode | Positive |
| Detector m/z Optimization | Low m/z |
| Source DC Offset (V) | 21 |
| In-source Trapping (V) | Off |
| Ion Transfer Target m/z | Low m/z |
| Injection Flatapole DC (V) | 5 |
| Inter Flatapole Lens (V) | 4 |
| Bent Flatapole DC (V) | 2 |
| Transfer Multipole DC (V) | 0 |
| C-Trap Entrance Lens Inject (V) | 2 |
| Trapping gas pressure setting | 3 |
| **Nano ESI parameters** | |
| spray Voltage(kV) | 1-1.4 |
| Capillary Temp (C°) | 200 |
| S-lens RF level | 200 |

### Supplementary Figures

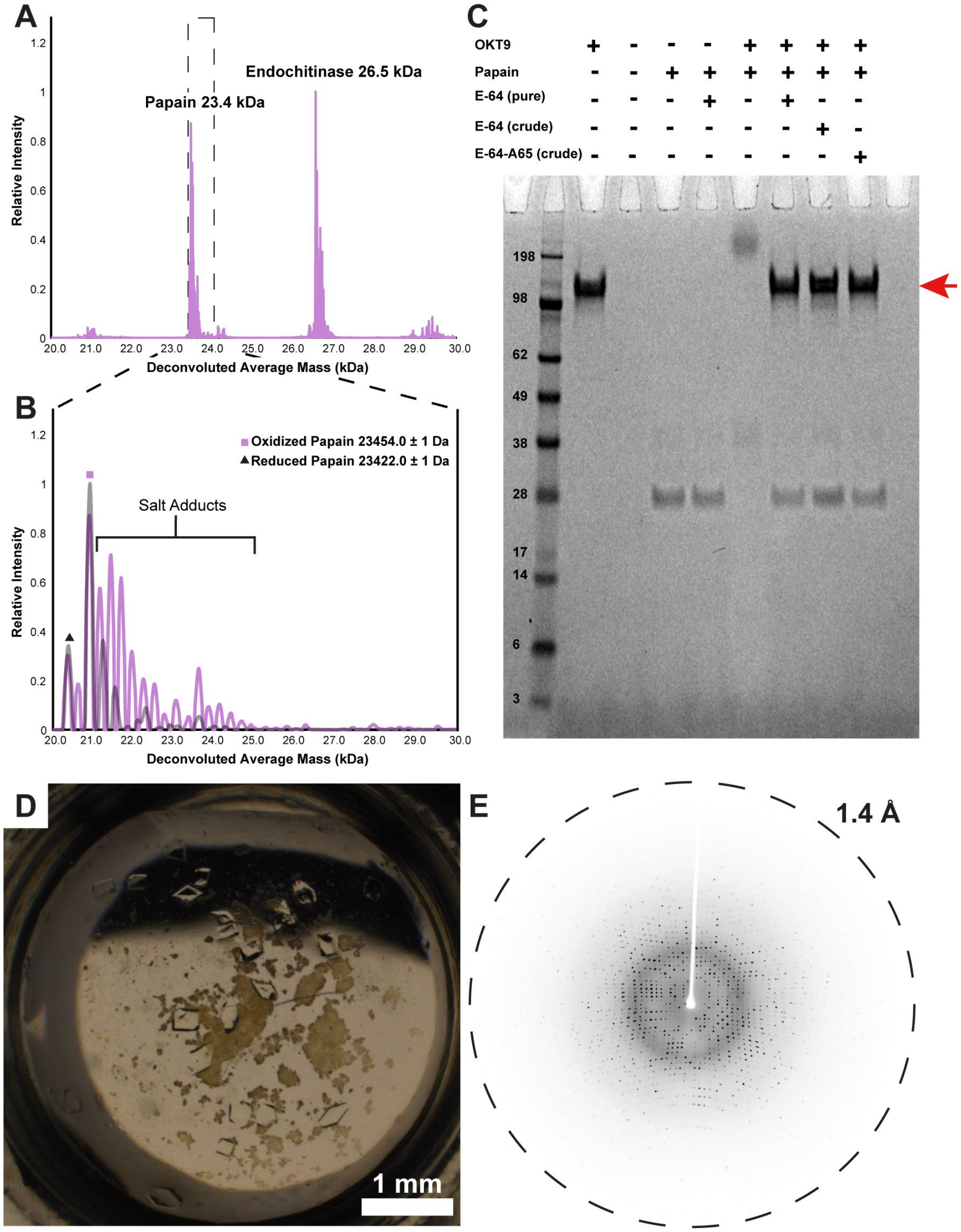

**Figure S1. Biochemical characterization of papain.** Native mass spectrum of twice-crystallized papain extract used for crystallography experiments (A), with enhanced view of the x-axis region of the spectrum encompassing the masses corresponding to papain present in the sample (B). SDS-PAGE gel indicating activity of papain present in the extract for cleaving model protein OKT9 (150 kDa) in the absence and presence of E-64, as well as crude in-house preparations of E-64 and the analog E-64-A65 from biosynthetic reaction (C). Representative crystals of papain from this sample (D) and single-crystal X-ray diffraction pattern of one such crystal (E).

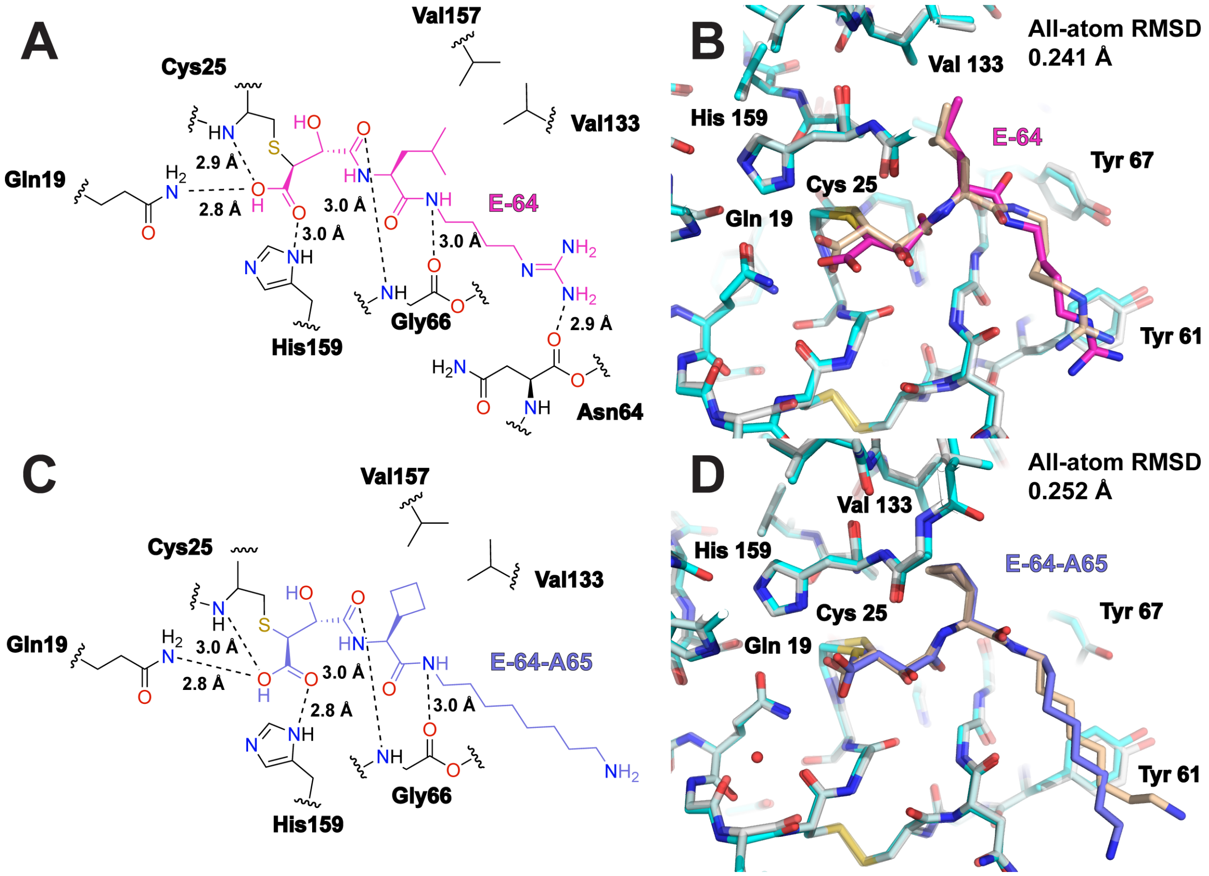

**Figure S2. Structure of papain active site complexed with E-64.** 2D representation of chemical structure of E-64 bound in papain's active site (A). Overlay of papain-E-64 active site determined by MicroED and the equivalent site determined by single crystal XRD (PDB ID: 9CKT) (B). 2D representation of chemical structure of E-64-A65 bound in papain's active site (C). Overlay of papain-E-64 active site determined by MicroED and the equivalent site determined by single crystal XRD (PDB ID: 9CKT) (D).

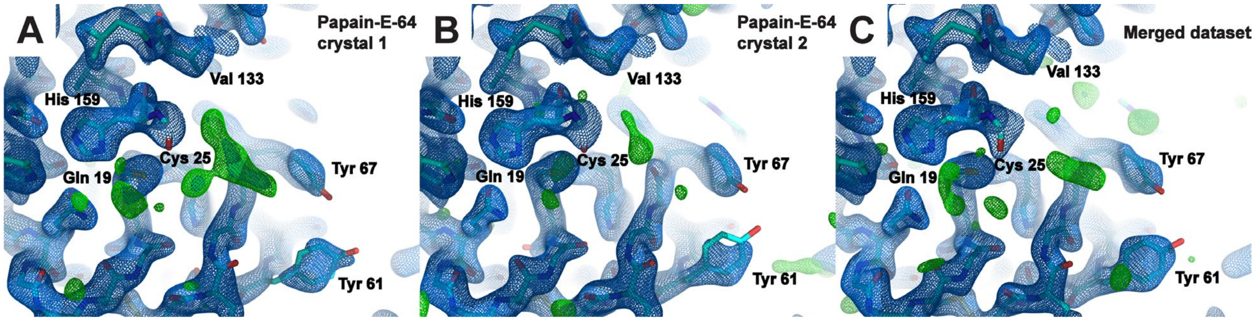

**Figure S3. Merging data from a few microcrystals may dilute ligand signal.** MicroED structure of the papain active site at 2.3 Å resolution from a single papain microcrystal co-crystallized with E-64 (A), and MicroED structure of the papain active site from a different papain microcrystal on the same TEM grid at 2.2 Å resolution (B) prior to modeling of any ligand in the active site. Density in the structure determined from the first crystal shows distinct evidence of an unmodeled ligand, while the structure determined from the second crystal shows only very weak unsatisfied density in the active site. When reflections from these two crystals are merged to yield a more complete dataset, the resulting structure has only weak residual density in the active site that might indicate a ligand's presence (C). Blue mesh indicates $2F_o$-$F_c$ map at 1.5σ levels and green mesh indicates $F_o$-$F_c$ map at 3σ levels.

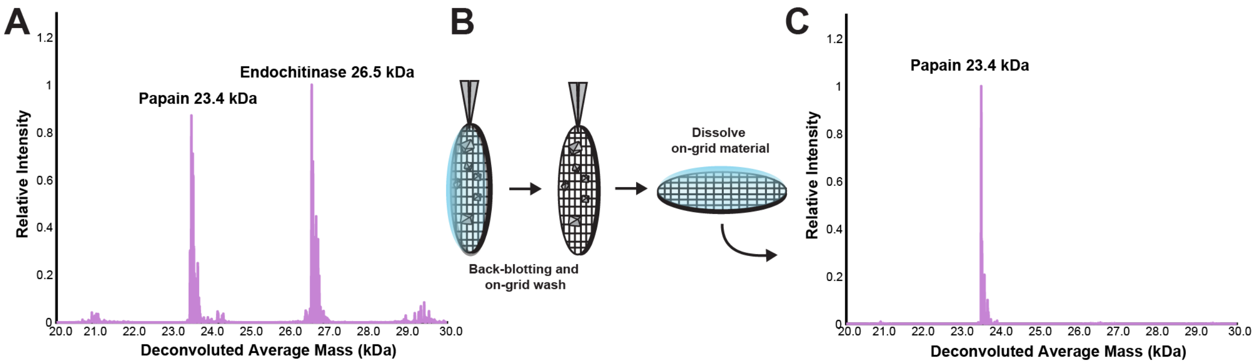

**Figure S4. Native MS analysis of MicroED samples.** Representative mass spectrum of papain extract in solution (A). Schematic of a mock ED-MS sample preparation, where an on-grid wash applied to the sample is successful at removing non-crystalline impurities prior to harvesting, dissolution, and nMS data collection (B). Representative mass spectrum of apo-form papain from dissolved crystals harvested from a cryoEM grid (C).

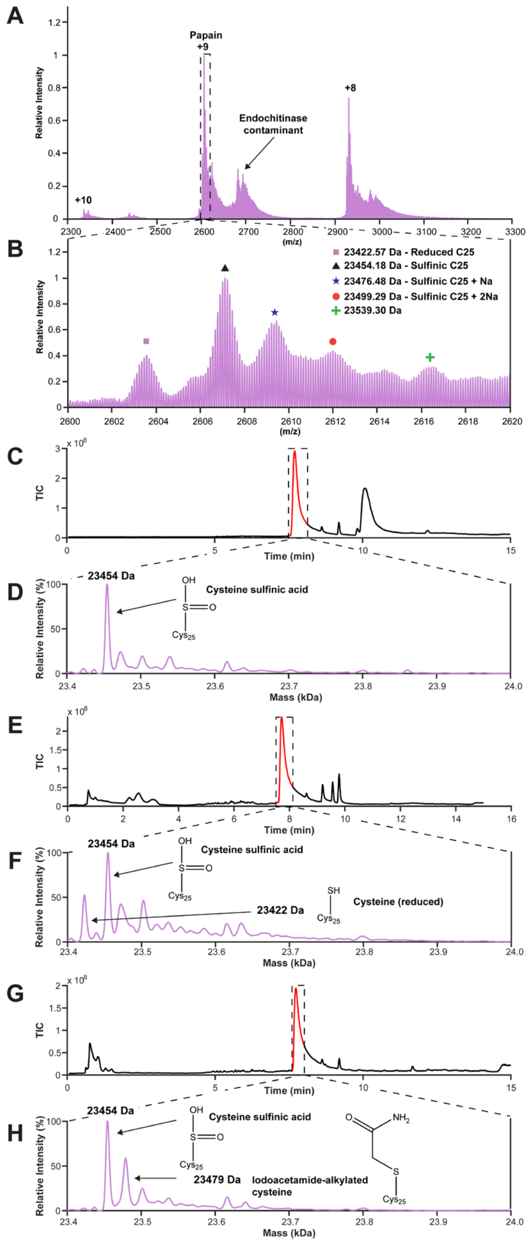

**Figure S5. LCMS analysis of Cys25 oxidation state in papain crystals.** High resolution native mass spectrum of twice-crystallized papain extract (A). Enlarged view of 2600 to 2620 m/z identifying average mass of isotopically resolved peaks in the +9 charge state of papain (B). Liquid chromatography plot and mass spectrum of the primary LC peak for papain without reducing agent present (C-D), papain in the presence of an excess of TCEP (E-F) and papain in the presence of an excess TCEP and the free cysteine binder iodoacetamide (G-H). Without reducing agent present, the sulfinic acid form of papain's catalytic cysteine 25 dominates (D), while application of reducing conditions using TCEP is able to recover some of the reduced cysteine form (F). Incubation of papain in reducing conditions with iodoacetamide confirms that the papain-iodoacetamide complex forms, indicated by a 57 Da mass-shifted peak in the mass spectrum relative to reduced papain (H). This indicates that in its reduced form Cys25 is available for binding to covalent ligands.

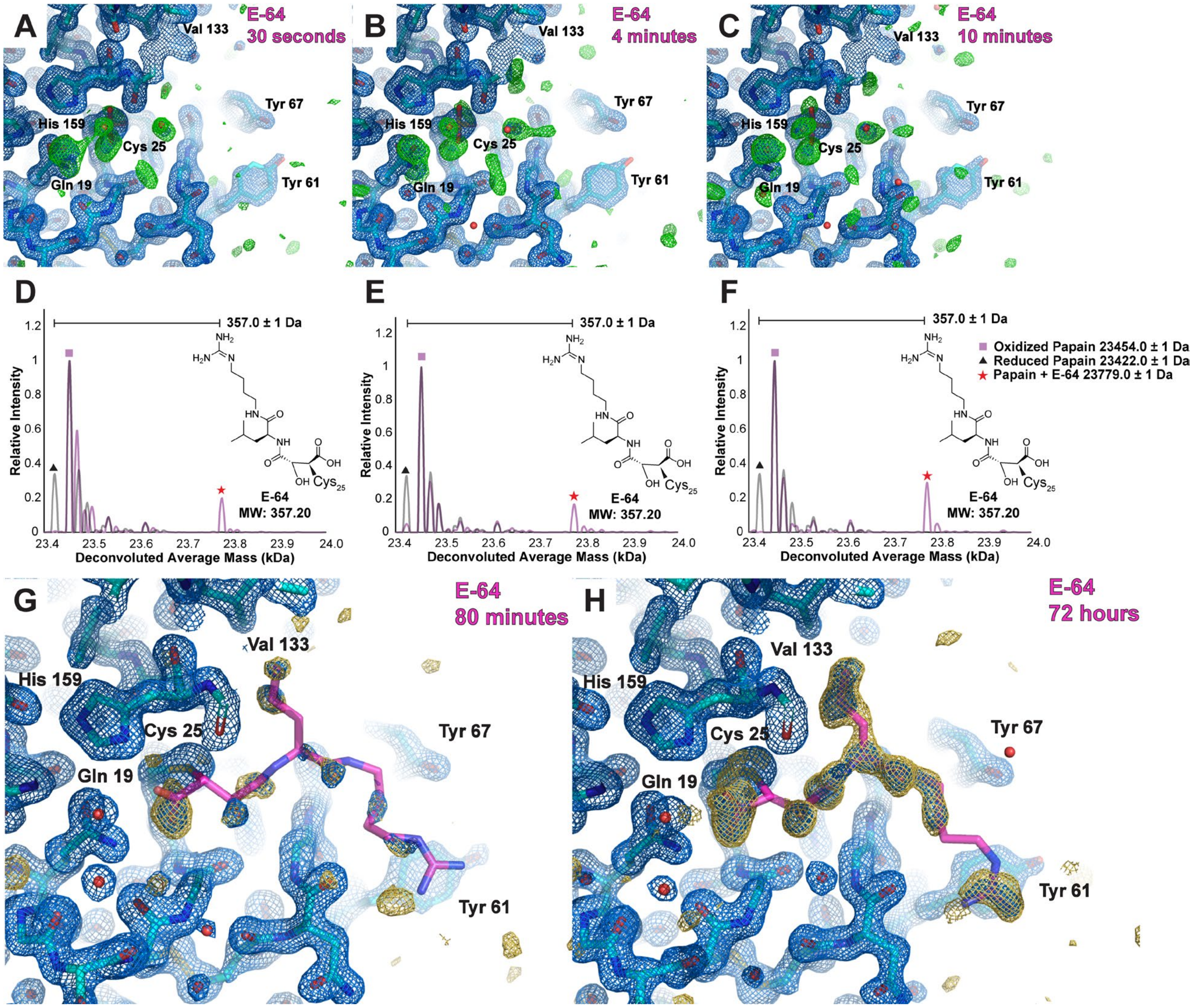

**Figure S6. Single crystal XRD of macroscopic papain crystals soaked with E-64 as a function of time.** scXRD structures of papain active site at 1.5 Å resolution from crystals frozen following 30 seconds (A), 4 minutes (B), and 10 minutes (C) of soaking with ~2.5 mM E-64. In each case, active site density corresponding to E-64 is not yet visible. Accompanying native mass spectrum measured from a dissolved macroscopic crystal treated in each way (D-F) reveals the presence of some papain-E-64 complex, in addition to the reduced cysteine and cysteine sulfinic acid apo-forms of papain, in each crystal sample, supporting that binding occurs but is diffusion-limited, and that the ligand requires additional time to permeate and bind throughout a macroscopic crystal. scXRD structures of papain active site from crystals frozen following significantly longer times soaking in an equivalent E-64 solution, for 80 minutes (structure at 1.6 Å resolution) (G) and 72 hours (structure at 1.5 Å resolution) (H), where evidence of the ligand becomes more apparent. Blue mesh indicates $2F_o$-$F_c$ map at 1.5σ levels, green mesh indicates $F_o$-$F_c$ map at 3σ levels at the current stage of refinement the figure at which the structure in the image is displayed, and gold mesh indicates the $F_o$-$F_c$ map at 3σ levels that was present prior to modeling a ligand which is ultimately satisfied once refinement is complete.

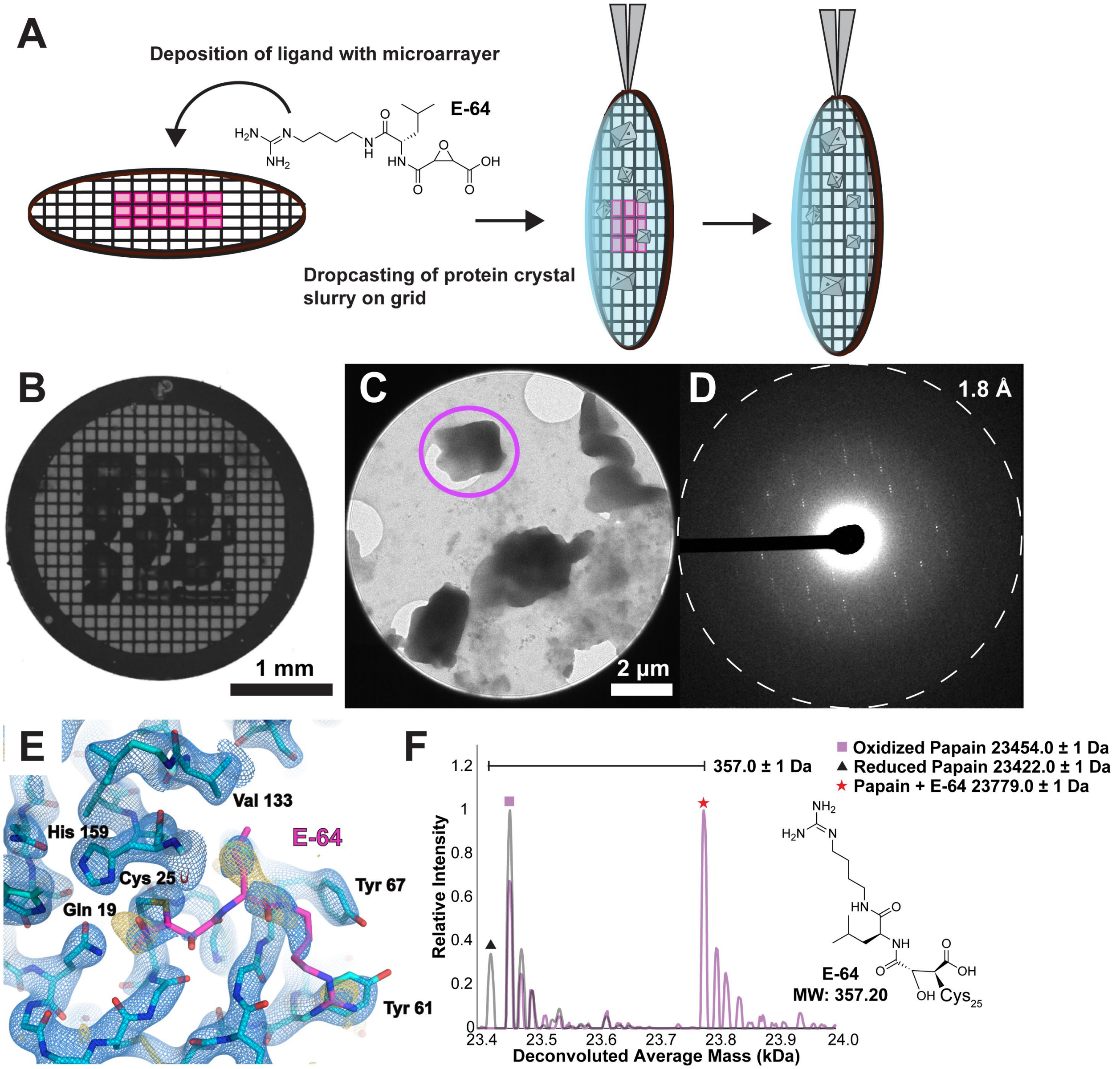

**Figure S7. MicroED structure determination of papain microcrystals prepared on ligand-arrayed TEM grids.** Schematic of small molecule printing on TEM grids with a microarrayer, after which protein microcrystals were incubated on the printed grid and incubated prior to blotting and plunge-freezing (A). Representative light microscopy image of one such grid following arraying with a 26 mM solution of E-64 (B). Representative microcrystal image (C) and electron diffraction pattern (D) of papain crystals prepared on arrayed grids in this fashion. 2.8 Å resolution MicroED structure of the papain active site from microcrystals mixed with arrayed E-64 ligand on grid, with $2F_o$-$F_c$ map in blue at 1.5σ following modeling of E-64 (magenta coordinates), alongside gold mesh indicating the $F_o$-$F_c$ map at 3σ levels that was present prior to modeling a ligand (E). nMS spectrum of grid-adsorbed material following MicroED collection, revealing presence of the papain-E-64 complex (F).

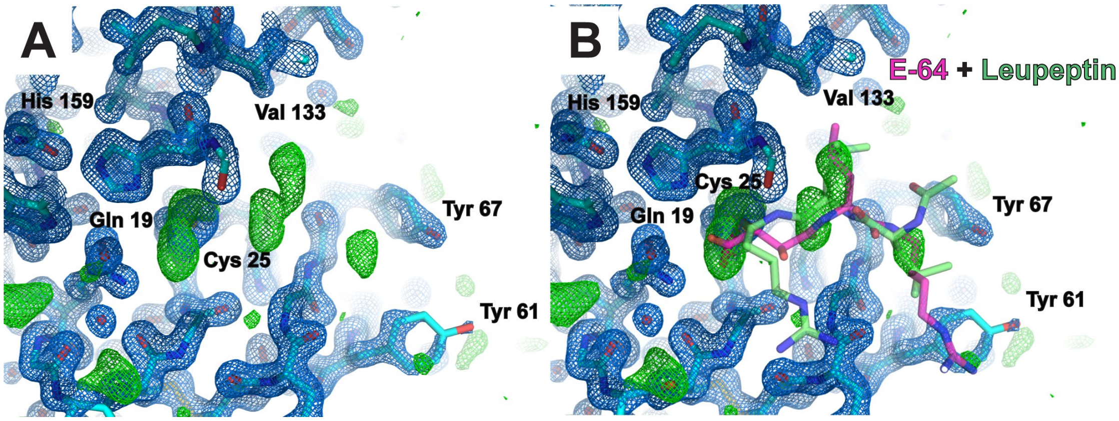

**Figure S8. Single crystal XRD structure of papain crystals soaked with a commercial protease inhibitor cocktail for 20 hours.** scXRD structure of the papain active site at 1.5 Å resolution from crystals frozen after 20 hours of soaking with commercially available protease inhibitor cocktail containing E-64, AEBSF, leupeptin, aprotinin, bestatin, and EDTA at varying relative concentrations. Model superimposed with $2F_o$-$F_c$ map at 1.5σ and $F_o$-$F_c$ map at 3σ following refinement without any ligand modeled (A), and the same structure overlayed with E-64 (magenta coordinates) and leupeptin (lime coordinates) in the active site (B). Either potential binder satisfies the active site density reasonably well, signaling that even with the improved resolution afforded by XRD compared to MicroED unambiguously distinguishing structurally similar ligands is non-trivial without nMS.

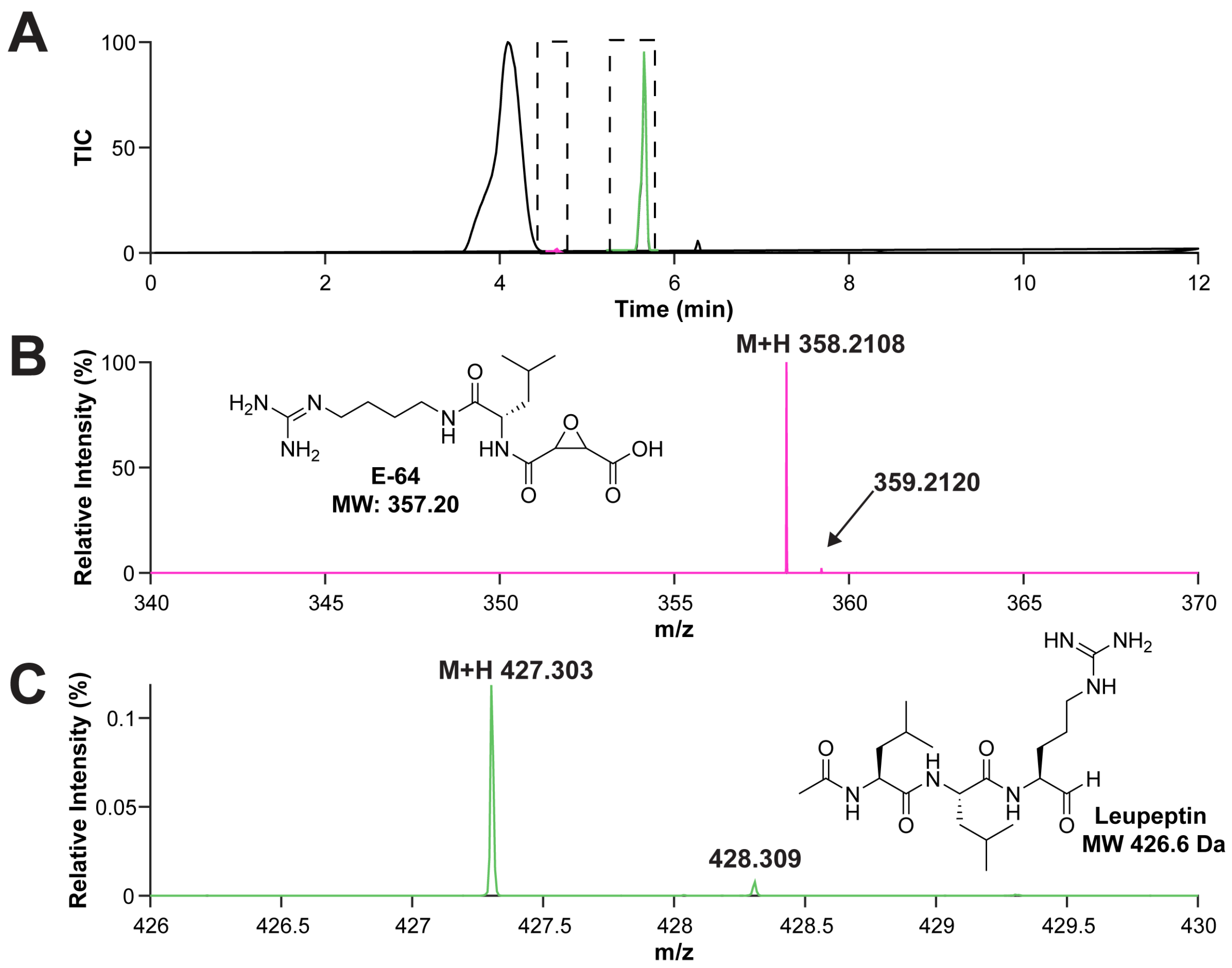

**Figure S9. LCMS analysis of commercial protease inhibitor cocktail.** Liquid chromatography plot from LCMS of protease inhibitor cocktail, with the elution peak for E-64 highlighted in magenta and elution peak for leupeptin highlighted in lime (A). Mass spectrum of the magenta LC peak in panel A confirming presence of E-64 (B) and mass spectrum of the lime LC peak in panel A confirming presence of leupeptin (C).

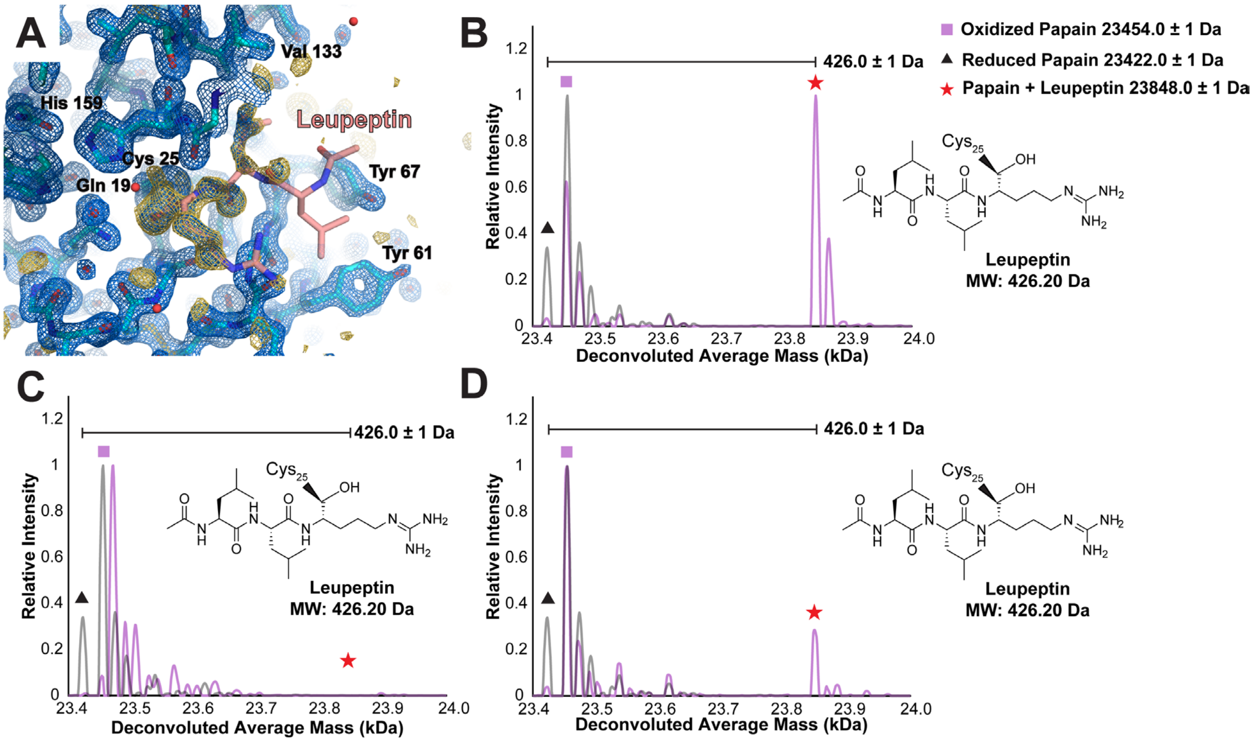

**Figure S10. Structures of the papain-leupeptin complex.** Single-crystal XRD structure at 1.6 Å resolution of papain co-crystallized with leupeptin with $2F_o$-$F_c$ map in blue at 1.5σ following modeling of leupeptin, alongside gold mesh indicating the $F_o$-$F_c$ map at 3σ levels that was present prior to modeling a ligand (A). Native mass spectrum of papain mixed with leupeptin in solution (B), macroscopic papain-leupeptin co-crystals suitable for single-crystal X-ray diffraction (C), and papain-leupeptin (micro) cocrystals harvested from a TEM grid (D).

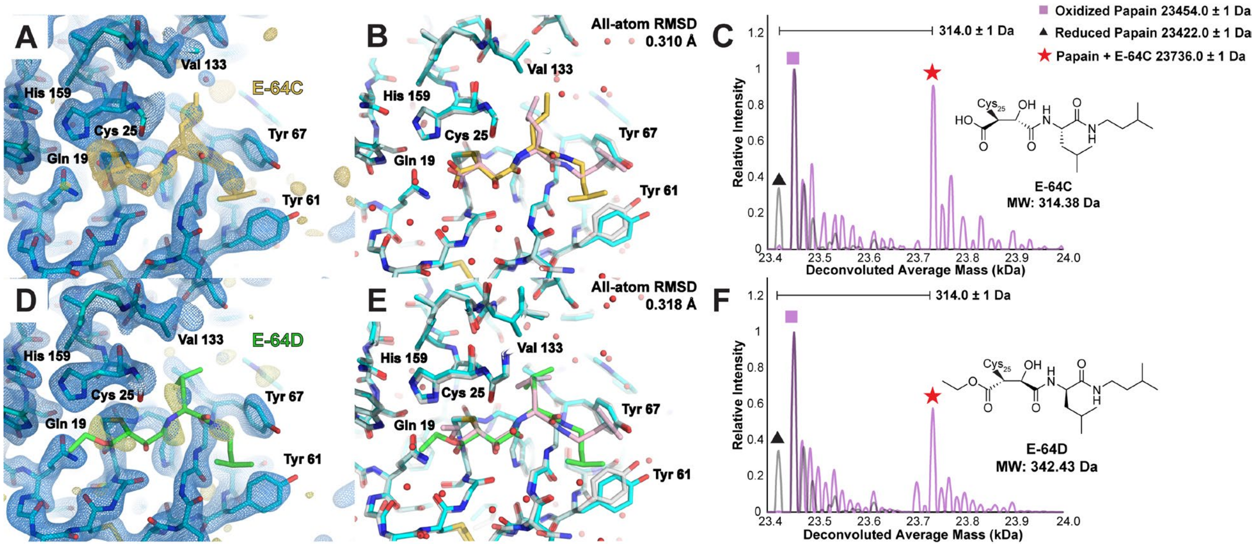

**Figure S11. MicroED structures of papain-E-64C and papain-E-64D cocrystals.** 2.2 Å resolution MicroED structure of papain co-crystallized with E-64C (A), and overlay of this structure with a single crystal XRD structure of the same complex (PDB ID: 9EG7) (B). Native mass spectrum of papain-E-64C co-crystals harvested from TEM grid after MicroED data collection (C). 2.3 Å resolution MicroED structure of papain co-crystallized with E-64D (D), and overlay of this structure with a single crystal XRD structure of the same complex (PDB ID: 9CKW) (E). Native mass spectrum of papain-E-64D co-crystals harvested from TEM grid after MicroED data collection (F). For structure images, the $2F_o$-$F_c$ map is shown in blue at 1.5σ following modeling of the ligand, alongside gold mesh indicating the $F_o$-$F_c$ map at 3σ levels that was present prior to modeling the ligand.

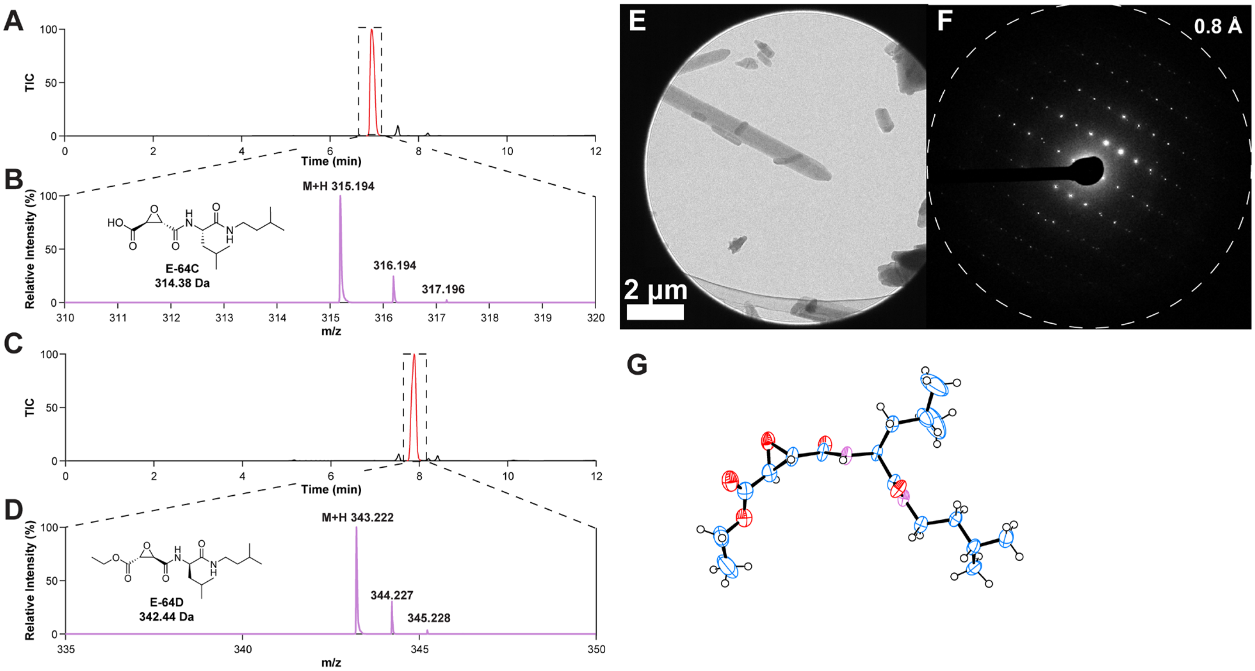

**Figure S12. Characterization of E-64D prior to soaking into crystals.** LCMS analysis of E-64C (A-B) and E-64D (C-D) in solution, revealing masses at the expected molecular weights for each. MicroED structure determination of E-64D powder, where nanocrystals (E) yielded single-crystal diffraction to high resolution (F). ORTEP diagram of MicroED structure of E-64 at 0.8 Å resolution, with the terminal ester bound to the epoxide warhead intact (G).

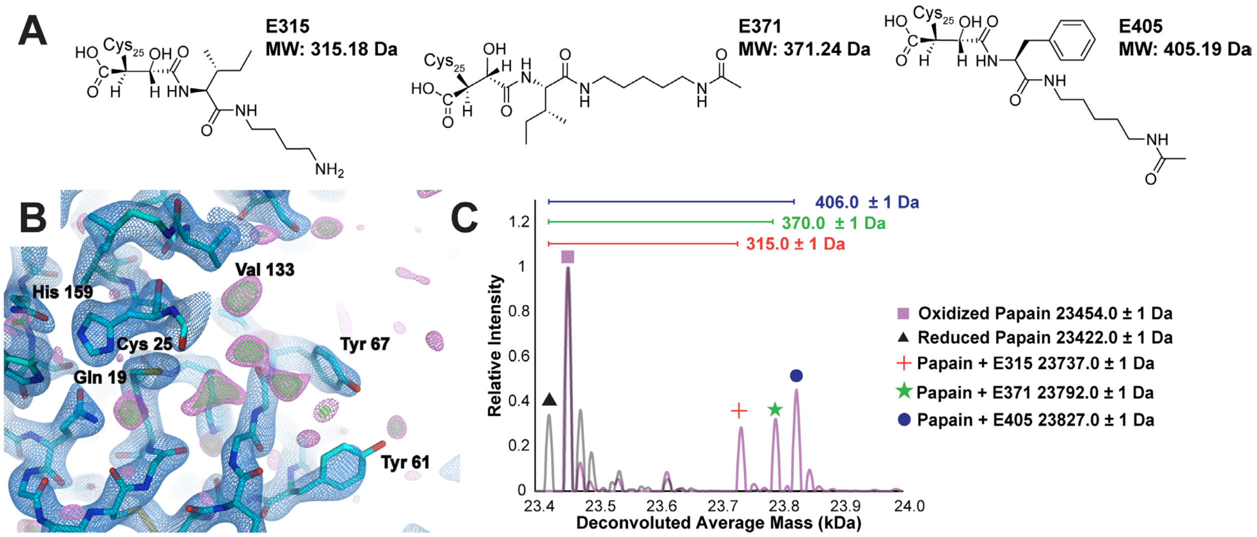

**Figure S13. ED+MS analysis of microcrystals soaked with an equimolar mixture of biosynthetic E-64 analogs without E-64-A65.** Upon noting by ED-MS that the natural product E-64-A65 bound papain and could be visualized in a MicroED structure, and was likewise detectable as part of a bound species within papain crystals soaked with a mixture of natural product E-64 analogs, a follow-up experiment was performed soaking papain crystals with the same natural product mixture excluding E-64-A65 (A) to identify weaker binders. 2.5 Å resolution MicroED structure of the papain active site soaked with an equimolar mixture of analogs E316, E372, and E406 for 10 minutes revealing weak residual density in the $F_o$-$F_c$ map within the active site, with $2F_o$-$F_c$ map at 1.5σ in blue, $F_o$-$F_c$ map at 3σ in green, and $F_o$-$F_c$ map at 2.5σ in magenta (B). Native mass spectrum measured from the same crystals harvested after MicroED data collection (C). While ligand density in the MicroED structure is too weak to easily interpret, nMS indicates that papain-E315, papain-E371, and papain-E405 are all present as bound species.

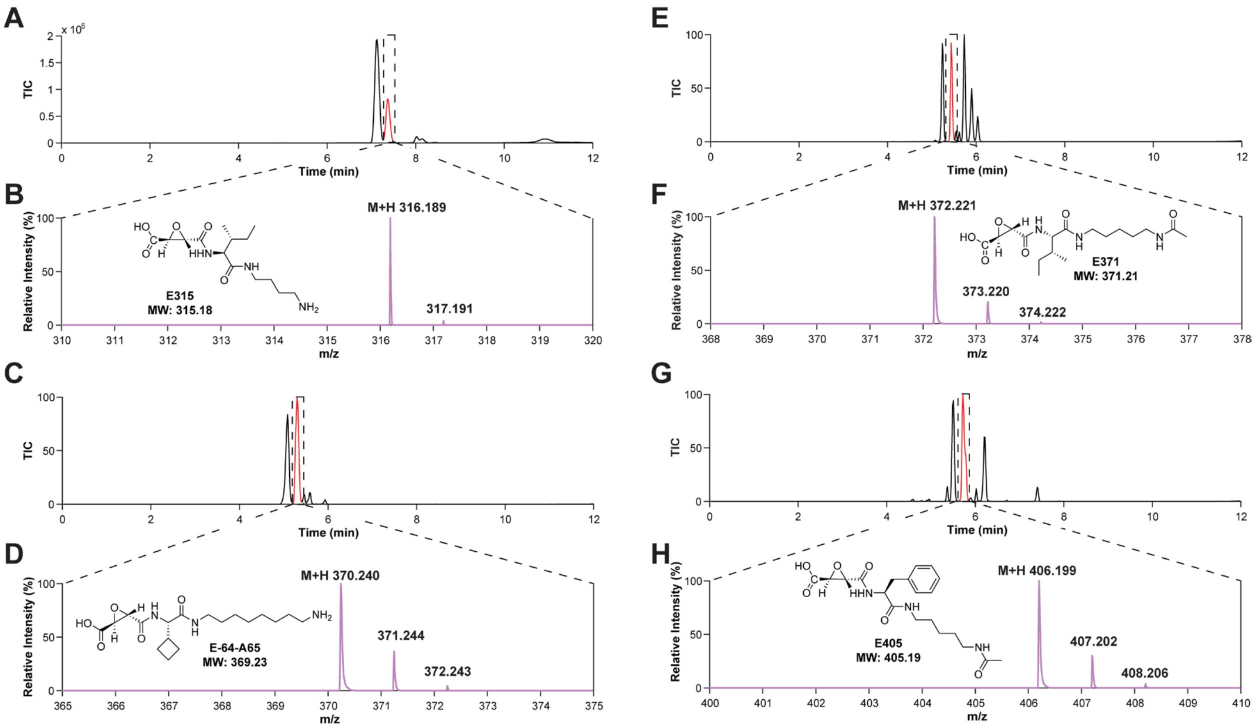

**Figure S14. LCMS of biosynthetic E-64 analogs.** LC-MS analysis of the biosynthetic E-64 analogs E315 (A-B), E-64-A65 (C-D), E371 (E-F), and E405 (G-H). Panels (A, C, E, G) display chromatograms with highlighted retention times indicating where mass spectra were extracted. Panels (B, D, F, H) present the corresponding mass spectra, confirming the molecular ion peaks for each analog.

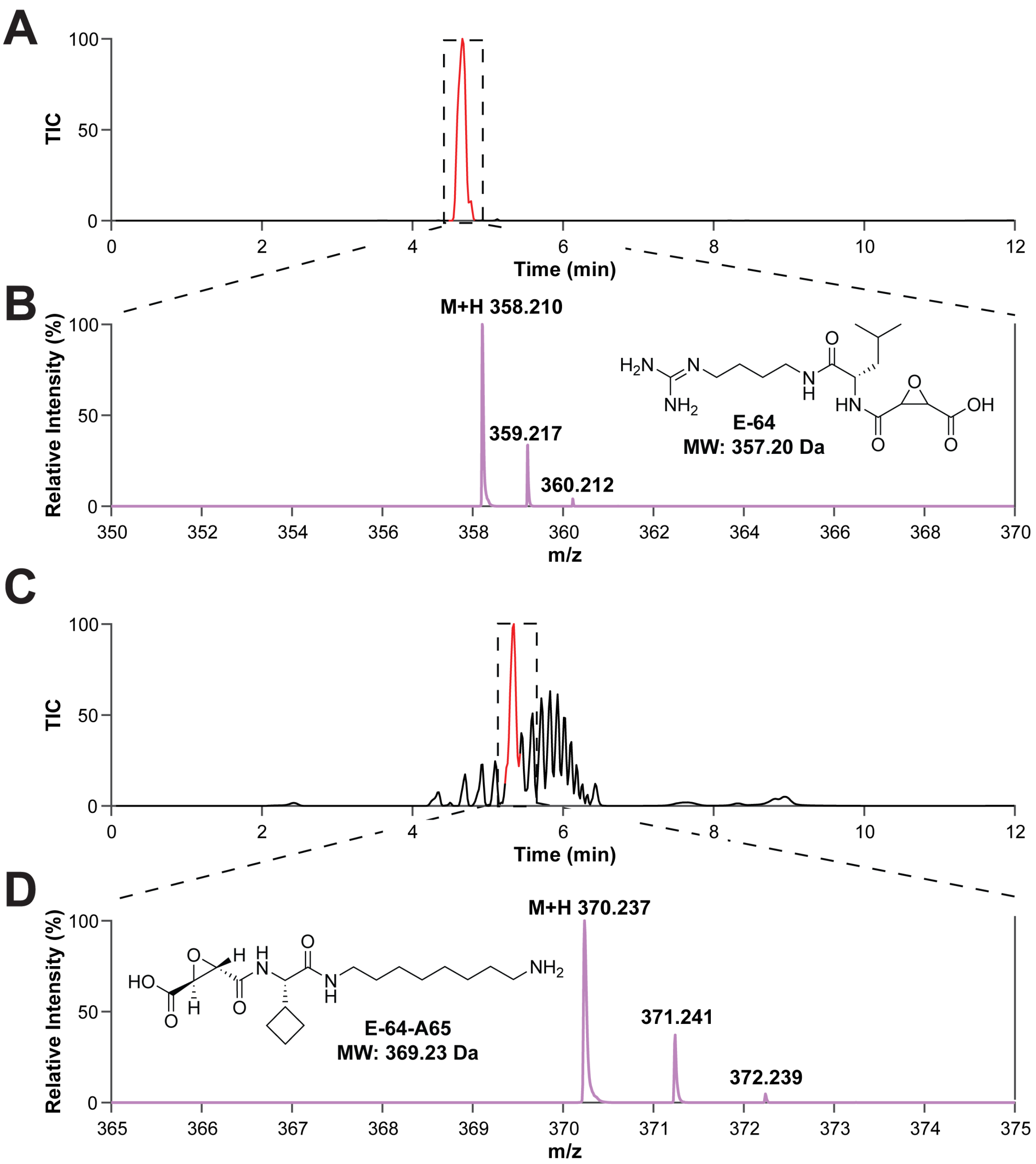

**Figure S15. LCMS analysis of crude biosynthetic products: E-64 and E-64-A65.** LC-MS analysis of E-64 (A-B) and E-64-A65 (C-D). Panels (A, C) display chromatograms with highlighted retention times indicating where mass spectra were extracted. Panels (B, D) present the corresponding mass spectra, confirming the molecular ion peaks for each compound.

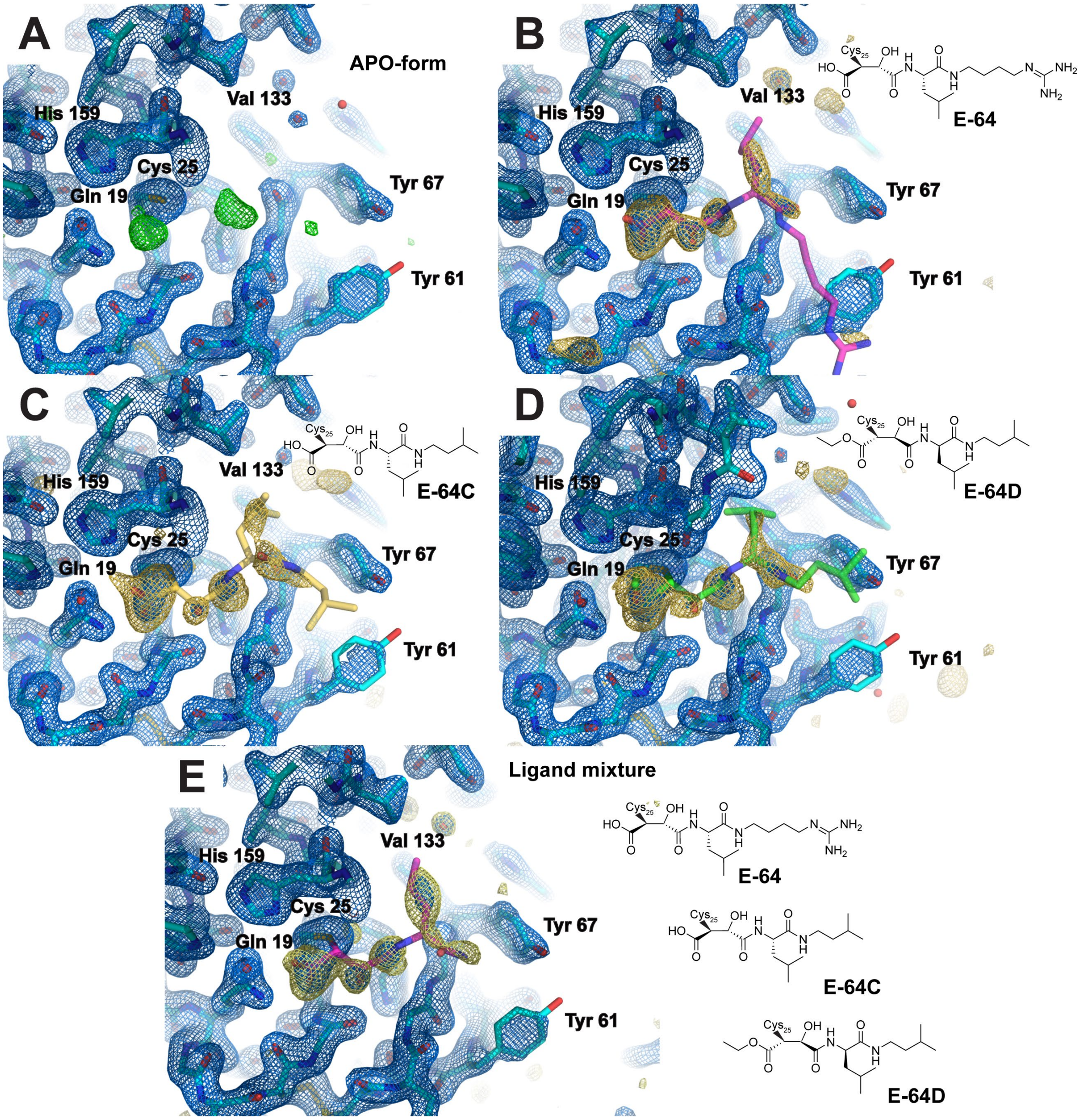

**Figure S16. Serial synchrotron XRD structures of papain and high-affinity ligands.** Active-site views at 1.8 Å resolution of papain microcrystal structures determined by serial synchrotron X-ray crystallography at beamline ID29 at the European Synchrotron Radiation Facility (ESRF) in apo-form (A), soaked with E-64 (B), soaked with E-64C (C), soaked with E-64D (D), and soaked with an equimolar mixture of all three ligands (E). For the structure determined from the mixture-soaking experiment, the portion of the ligand that is structurally conserved across all three potential ligands can be modeled into the difference density. Blue mesh indicates $2F_o$-$F_c$ map at 1.5σ levels, green mesh indicates $F_o$-$F_c$ map at 3σ levels at the current stage of refinement the figure at which the structure in the image is displayed, and gold mesh indicates the $F_o$-$F_c$ map at 3σ levels that was present prior to modeling a ligand which is ultimately satisfied once refinement is complete.

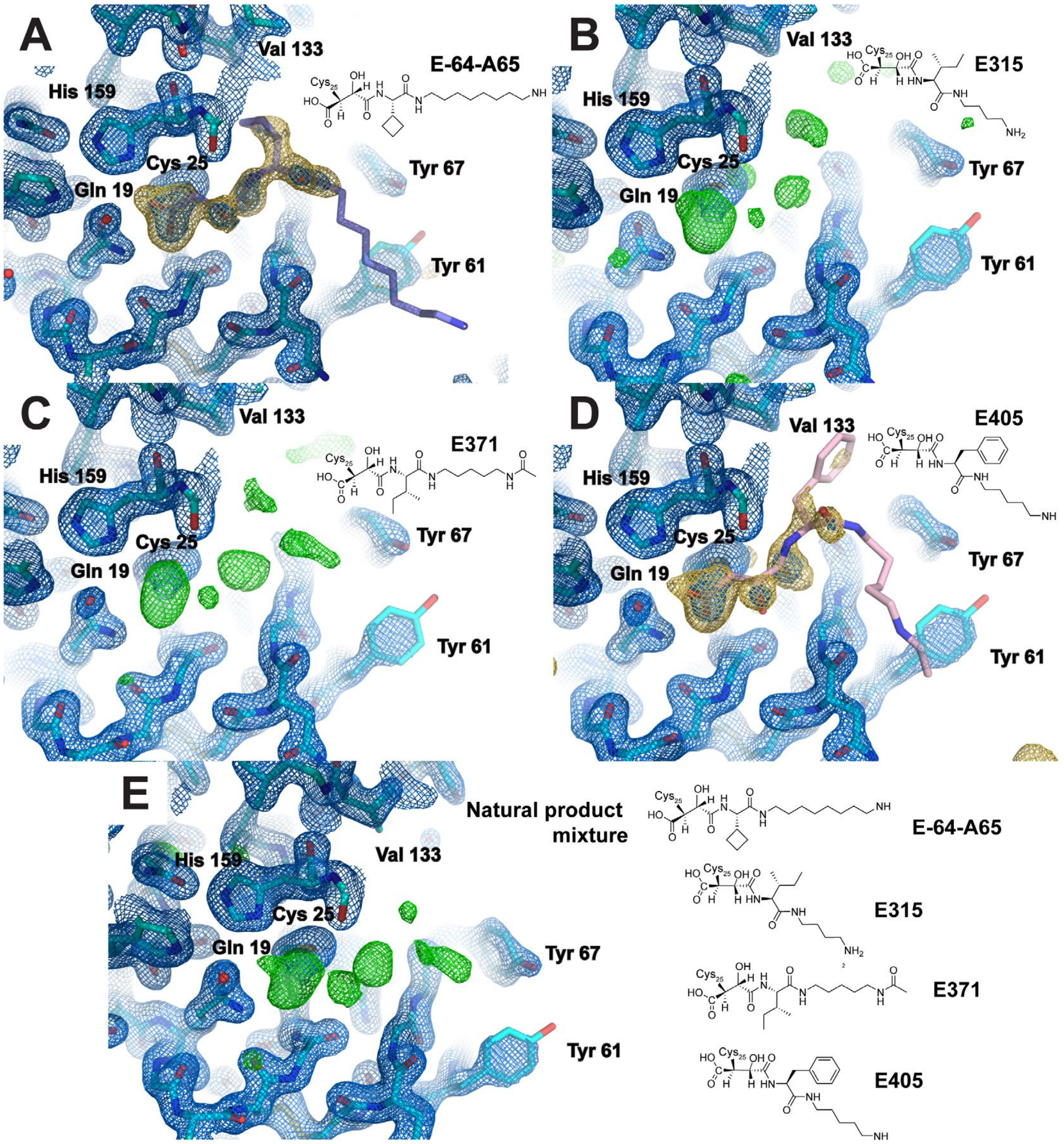

**Figure S17. Serial synchrotron XRD structures of papain soaked with natural product inhibitors.** Active-site visualizations at 1.8 Å resolution of structures from papain microcrystals determined by serial synchrotron X-ray diffraction at Beamline ID29 at the European Synchrotron Radiation Facility (ESRF). Microcrystals were soaked with either E-64-A65 (A), E315 (B), E371 (C), E405 (D) or an equimolar mixture of all four (E). In cases where the ligand density was clear enough to interpret, as was true for E-64-A65 (panel A, violet coordinates) and E405 (panel D, pink coordinates), the ligand was modeled in the active site and the structure refined. Less well-defined density was measured from crystals soaked with E315 and E371, though the presence of strong peaks unaccounted for by the apo-form structure indicate that some trace binding of these ligands may have occurred. Blue mesh indicates $2F_o$-$F_c$ map at 1.5σ levels, green mesh indicates $F_o$-$F_c$ map at 3σ levels at the current stage of refinement the figure at which the structure in the image is displayed, and gold mesh indicates the $F_o$-$F_c$ map at 3σ levels that was present prior to modeling a ligand which is ultimately satisfied once refinement is complete.

**CheckCIF/Platon report for MicroED structure of E-64D**

Included are responses to A and B level alerts

### Alert level A

PLAT029_ALERT_3_A _diffrn_measured_fraction_theta_full value Low . 0.919 Why?

Complete sampling of reflections in MicroED is often limited by accessible tilt range of the TEM, especially in cases where crystals suffer from orientation bias as these thin, plate shaped microcrystals did.

### Alert level B

PLAT082_ALERT_2_B High R1 Value .................................. 0.16 Report
PLAT084_ALERT_3_B High wR2 Value (i.e. > 0.25) ................... 0.41 Report

Greater refinement R factors are anticipated in MicroED, when compared to XRD. These mirror the greater R-merge values enountered in data reduction, which are known to be inflated in MicroED by inelastic scattering, dynamical scattering, and other unmodeled aberrations.

PLAT340_ALERT_3_B Low Bond Precision on C-C Bonds ............... 0.01954 Ang.

Limited completeness in MicroED data may result in resolution anisotropy of the final electrostatic potential map. This may reduce bond length precision, and might be mitigated by application of further restraints during refinement, at the expense of R-factors.

```
PLAT911_ALERT_3_B Missing FCF Refl Between Thmin & STh/L=   0.600      158 Report
           0  2  0,   0  4  0,   1  5  0,   0  6  0,   2  6  0,   2  7  0,
           0  8  0,   0 10  0,   1 10  0,   1 11  0,   0 12  0,   1 12  0,
           1 13  0,   0 14  0,   1 14  0,   1 15  0,   0  2  1,   0  3  1,
           0  4  1,   0  5  1,   0  6  1,   0  7  1,   0  8  1,   0  9  1,
           0 10  1,   0 11  1,   1 11  1,   0 12  1,   1 12  1,   0 13  1,
           1 13  1,   0 14  1,   1 14  1,   0 15  1,   1 15  1,   0  3  2,
           0  4  2,   0  5  2,   0  6  2,   0  7  2,   0  8  2,   0  9  2,
           0 10  2,   0 11  2,   0 12  2,   0 13  2,   1 13  2,   0 14  2,
           1 14  2,   0 15  2,   1 15  2,   0  4  3,   0  5  3,   0  6  3,
           0  7  3,   0  8  3,   0  9  3,   0 10  3,   0 11  3,   0 12  3,
           0 13  3,   0 14  3,   1 14  3,   0 15  3,   1 15  3,   0  0  4,
           0  5  4,   0  6  4,   0  7  4,   1  7  4,   0  8  4,   0  9  4,
           0 10  4,   0 11  4,   0 12  4,   0 13  4,   0 14  4,   0 15  4,
           1 15  4,   0  6  5,   0  7  5,   0  8  5,   0  9  5,   0 10  5,
           0 11  5,   0 12  5,   0 13  5,   0 14  5,   0 15  5,   0  0  6,
           0  4  6,   0  7  6,   0  8  6,   0  9  6,   0 10  6,   0 11  6,
```

See response to PLAT082 alert.
